# Biphasic release propensity of mucin granules is supervised by TSPAN8

**DOI:** 10.1101/2022.09.14.507971

**Authors:** Wojnacki José, Lujan Agustín, Foresti Ombretta, Aranda Carla, Bigliani Gonzalo, Maria Pena Rodriguez, Brouwers Nathalie, Malhotra Vivek

## Abstract

Agonist-mediated stimulated pathway of mucin and insulin release is biphasic in which a rapid fusion of pre-docked granules is followed by slow docking and fusion of granules from the reserve pool. The sustained neurotransmitter release also necessitates docking of vesicles from a reserve pool. We present here a surprising finding that plasma membrane-located tetraspanin-8 (Tspan-8) sequesters syntaxin-2 (Stx2) to control external agonist-dependent mucin release. Tspan-8 specifically affects fusion of granules in reserve during the second phase of stimulated mucin release. The Tspan-8 and Stx2 complex does not contain VAMP-8 and Munc18, which are required for fusion of mucin granules. We suggest that by sequestering Stx2, Tspan-8 prevents docking granules in the reserve pool. In the absence of Tspan-8, granules in reserve pool are free to dock to Stx2 and their fusion doubles the quantities of mucins secreted. Tspan-8 thus emerges as the long-sought component that controls biphasic mucin release. We suggest a similar mechanism likely controls biphasic insulin and sustained neurotransmitter release.

## Introduction

A key issue in cell and tissue function is how cells secrete the right quantities of proteins. This becomes especially important for proteins such as neurotransmitters, insulin, and mucins. Cells have developed systems that allow proteins to be secreted at a slow but constant rate and also, an agonist-dependent stimulated release. When an external agonist binds to specific receptors, it causes an increase in intracellular calcium that triggers the fusion of large granules containing insulin or mucins (1-3) or in neurons, small synaptic vesicles that store neurotransmitters (4). Humans express 5 gel-forming mucins that are secreted by specialized goblet cells (5-7). Secreted mucins mix with other extracellular components to constitute the mucus, which acts as a lubricant and a protective barrier to the underlying epithelium (8-10). But, as noted in human pathologies of the airways and the digestive system (11-13), too much or too little extracellular mucins may be damaging for the underlying tissue, which begs the question, how is the amount of mucins secreted controlled by goblet cells?

Mature mucin granules fuse to the plasma membrane at a constant low rate (basal secretion) and, if necessary, an external agonist triggers a massive release of mucins in a short period of time (stimulated secretion) (14-16). Stimulated secretion is biphasic. Within seconds after agonist stimulation the rate of mucin secretion increases more than a thousand times compared to baseline secretion. The short burst of secretion (less than 1 minute) is followed by a longer, slower-rate phase. The second phase lasts several minutes during which, the secretion rate is 47 times slower compared to the peak rate but still 38 times higher compared to baseline secretion (14). It is generally assumed that the rapid release involves fusion of pre-docked vesicles, whereas the slower-sustained release involves fusion of vesicles/granules in reserve. The availability of specific sites, to which vesicles can dock, might be the major bottleneck controlling fusion (17), but how this is regulated remains unexplored. The highly conserved, SNARE proteins are essential for basal and stimulated granule fusion (4, 15, 18, 19). Mucin granules bearing the R-SNARE, Vesicle-associated membrane protein 8 (VAMP-8), fuse to the plasma membrane to release their contents (18, 20). During exocytosis, VAMP proteins interact with syntaxins (Q-SNAREs) in trans, to generate the necessary force for membrane fusion (4, 21, 22). Cells from the human airways express syntaxins 1, 2, 3 and 4 (23-25)but the identity of the syntaxin required for mucin secretion in intact cells remains unknown. Syntaxins are also required to dock secretory granules to the plasma membrane (26-28). It has been suggested that a release site (vesicle docking site) can exist in three different states: (i) empty and accessible for a vesicle (ii) occupied, ready for its vesicle to exocytose (iii) empty and refractory (not accessible for a vesicle) (17). Cells could control the availability of Q-SNAREs at the plasma membrane to increase or decrease the number of accessible docking sites and the quantity of released material during secretion.

We report here that syntaxin-2 (Stx2) is present on the apical plasma membrane of mucin-secreting cells and is required for mucin secretion. A more surprising finding is that tetraspanin-8 (Tspan-8), which also localizes to the plasma membrane, binds syntaxin-2 (and syntaxin-3) and controls its availability for SNARE-dependent mucin secretion. Stx2 is either bound to VAMP-8 or to Tspan-8. When Stx2 is in complex with Tspan-8, it’s unavailable to engage with VAMP-8. Loss of Tspan-8 increases mucin release by the stimulated pathway, specifically during the second phase of release. In the absence of Tspan-8, the biphasic release of mucins is deregulated. TSPAN8 thus emerges as component that control the release propensity of mucin granules by the external agonist stimulated pathway.

## Results

### Upregulation of Tetraspanin-8 in mucin secreting goblet cells

Upon differentiation, goblet cells upregulate mucin (MUC) genes and we asked a simple question. Are there genes, other than the mucins, that are upregulated during this event? A bioinformatics analysis of a single-cell RNA sequencing (scRNAseq) profiling database of healthy human airways (29) was thus performed. We compared the scRNAseq profiles of 17712 secretory cells with 24138 basal cells from four regions of the human airways: nasal, proximal, intermediate and distal (Figure Supplementary 1A). For each one of these regions we calculated the fold change in gene expression between secretory and basal cells. Most genes were repressed in differentiated secretory cells, which is shown as dots below the horizontal dashed line in the dot plot (Figure 1A and Supplementary 1B). Genes above the horizontal line were upregulated in secretory cells. As expected, MUCAC and MUC5B are highly upregulated. Of interest was the finding that TSPAN8 is upregulated (Figure 1A and Supplementary 1B). Neither the function of TSPAN8, nor its enrichment in goblet cells have been described. TSPAN8 expression in secretory cells from all areas of the airways was on average 42% higher than in basal cells and comparable to two secreted mucins, MUC5AC and MUC5B (Figure 1B). Less than 10% of basal cells expressed TSPAN8, MUC5AC or MUC5B while the same genes were expressed in over 75% of differentiated secretory cells. The proportion of basal, parabasal, multiciliated and secretory cells expressing ATG5 and GAPDH was similar in all cell types (Figure 1C). We also calculated the proportion of cells co-expressing TSPAN8, MUC5B and MUC5AC, and found that at least 60% of secretory cells co-expressed these proteins (Figure 1D).

**Figure 1.**
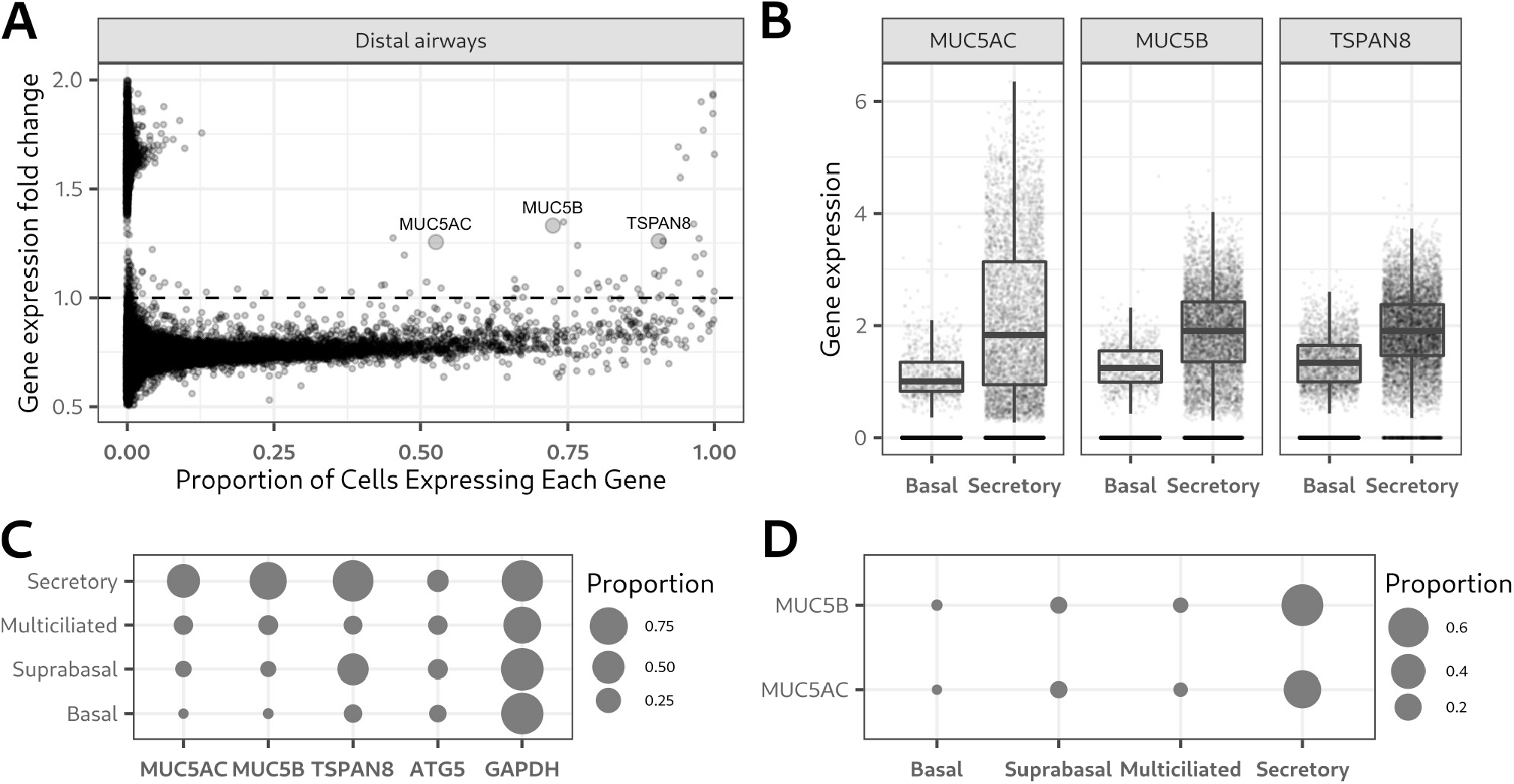
Tetraspanin-8 is upregulated in mucin-secreting cells from the healthy human airways. **A**. Dot plot showing fold-change in gene expression between basal and mucin-secreting cells in the y-axis and the proportion of mucin-secreting cells expressing each gene in the x-axis. Each dot represents one gene of the distal airways. Larger dots represent MUC5AC, MUC5B and TSPAN8 genes. Genes below the dashed horizontal line are down regulated in mucin secreting cells whereas genes above the line are up regulated. Dots are plotted with a 50% transparency value for better visibility in areas of the graph with high dots density. **B**. Box and dot plots of the expression levels of MUC5AC, MUC5B and TSPAN8 in basal and mucin-secreting cells from the human airways. Cells from the nasal, proximal, intermediate and distal regions of the airways were pooled together. Each dot represents gene expression in a single cell. Cells in which no expression was detected are lined at the bottom of the y-axis. The boxes of the box plot were calculated excluding cells with no detectable expression. **C**. Plot of the proportion of cells expressing MUC5AC, MUC5B, TSPAN8, and as control genes ATG5 and GAPDH. The proportion of cells expressing each gene was calculated for secretory, multiciliated, suprabasal, and basal cells. The area of the dot represents the proportion of cells expressing each gene according to the scale shown. **D**. Mucins and TSPAN8 co-expression plot. The proportion of cells co-expressing TSPAN8/MUC5AC and TSPAN8/MUC5B were calculated for secretory, multiciliated, suprabasal and basal cells. The area of the dot represents the proportion of gene co-expression according to the scale.

To test the role of TSPAN8 in mucin secretion, we used the human-derived and mucin-secreting cell line HT29-N2 (HT29 henceforth) (30, 31). A western blot analysis showed that HT29 cells express TSPAN8 and that its expression increases after differentiation (Figure Supplementary 1C and D).

### Tetraspanin-8 depletion causes mucin hyper-secretion

We first created a cell line that expresses endogenously tagged MUC5AC. We tagged the c-terminal tail of the MUC5AC locus with the super-folded Green Fluorescent Protein (sfGFP) (32) sequence by CRISPR/Cas9 genome editing. The GFP-tagged mucin-5AC-secreting cell line differentiated as wild type cells shown by the gradual increase in the levels of mucin-5AC during differentiation (Figure Supplementary 2A and B). Basal and ATP-stimulated mucin-5AC secretion were not affected in mucin-5AC·GFP-tagged cells (Figure Supplementary 2C and D; Supplementary Video 1). The total mucin-5AC content was not altered by addition of the sfGFP sequence (Figure Supplementary 2E). In differentiated cells, mucin-5AC·GFP localized to distinct granules that showed a very high degree of co-localization with the immunolabelled mucin-5AC (Figure Supplementary 2F; arrows). We also detected a minor population of GFP-positive granules, close to the perinuclear region, that were not recognized by the antibody for mucin-5AC suggesting that these could be immature mucin, without a mature epitope (Figure Supplementary 2F; arrowheads). Mucin-5AC·GFP did not co-localize with markers of the Golgi apparatus or with the lysosomal marker Lamp1 (Figure Supplementary 2G – I) suggesting that most GFP-positive granules are post-Golgi and that mucin-5AC·GFP is not en route to degradation. These data show that GFP-tagged and untagged mucin-5AC behave identically, and we could use this genetically-engineered cell line to visualize mucin-5AC localization and secretion.

We knocked-out (KO) TSPAN8 by CRISPR/Cas9 genome editing in the mucin-5AC GFP-tagged cell line. Complete loss of tetraspanin-8 mRNA and protein were confirmed by reverse transcription-polymerase chain reaction (RT-PCR) and western blot, respectively (Figure Supplementary 3A and B). A secretion assay showed that basal mucin-5AC·GFP secretion was unaltered, while ATP-stimulated secretion was increased by more than 50% in two independent TSPAN8 KO cell lines (Figure 2A and B). Equal amounts of cells in each condition was confirmed by western blot analysis of beta-tubulin in the cell lysates (Figure Supplementary 3C and D). Mucin-5AC·GFP expression was not changed by TSPAN8 KO (Figure 2C and D). Beta-tubulin in the secreted fractions was never above 3% of the total amount, statistically equal in WT and TSPAN8 KO cells, and unaffected by ATP stimulation (Figure Supplementary 3E and F). The increased mucin secretion upon TSPAN8 loss is therefore not due to altered gene expression or cell lysis.

**Figure 2.**
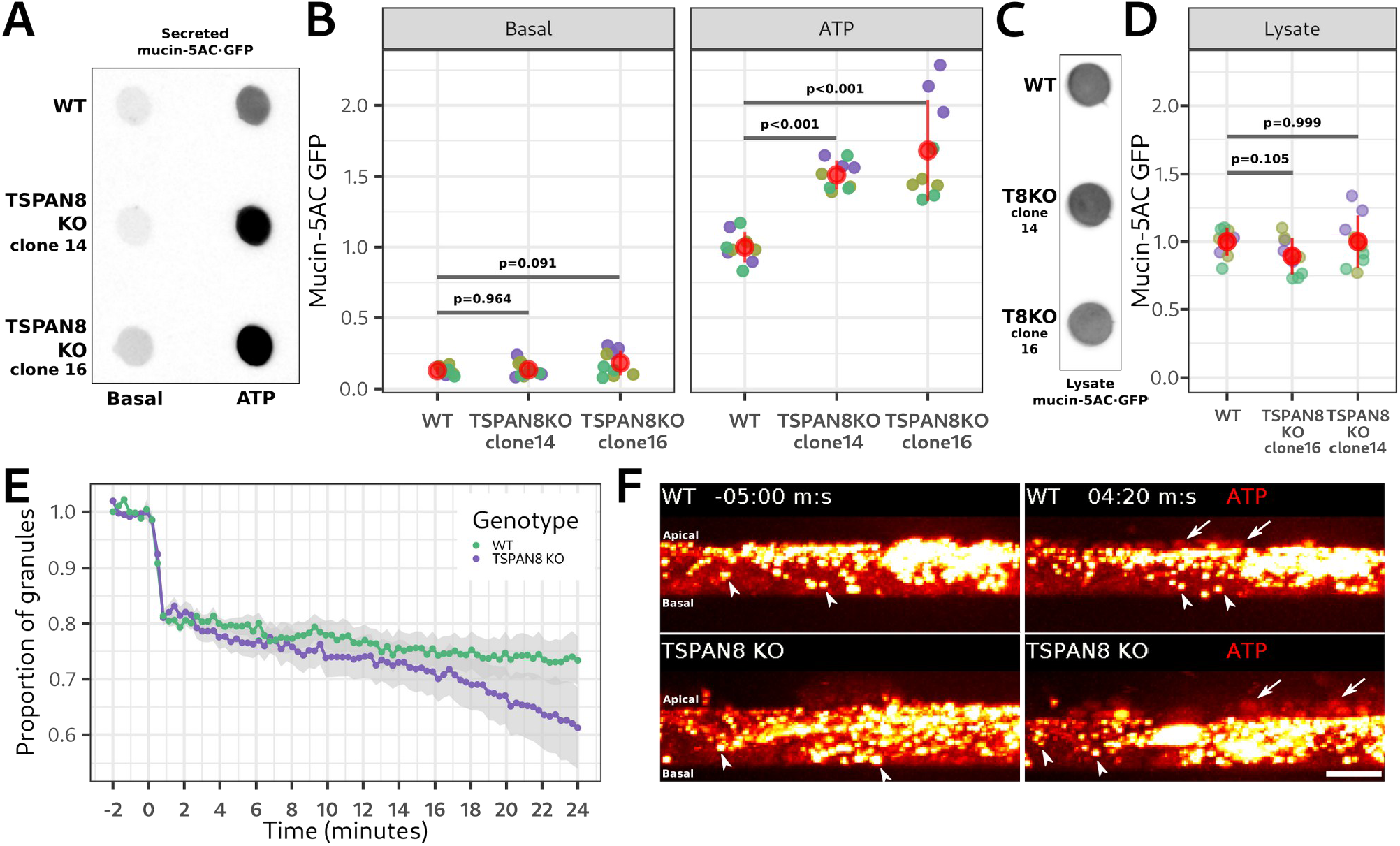
TSPAN8 KO increases mucin secretion. **A**. Representative dot blot showing mucin-5AC·GFP secreted by WT and TSPAN8 KO cell lines (clones 14 and 16). Left column is non stimulated (basal) secretion and right column is ATP-stimulated secretion. The fluorescent emission of sfGFP is shown. **B**. Quantification of the amount of mucin-5AC·GFP in the secreted fractions of basal and ATP-stimulated WT and TSPAN8 KO cell lines. A representative dot blot is shown in **A**. Each dot represents the signal from a single secretion assay. Grouped in different colors are secretion assays that were processed in parallel. Total number of replicates are 9. Red dots represent the mean +/- the standard deviation. Statistical analysis was performed independently for basal and ATP-stimulated conditions. The p values from a one-way ANOVA analysis with orthogonal contrasts are shown. **C**. Representative dot blot of the total (cell lysate) mucin-5AC·GFP in non stimulated WT and TSPAN8 KO cell lines. The fluorescent emission of sfGFP is shown. **D**. Quantification of the amount of mucin-5AC·GFP in the cell lysates of non stimulated WT and TSPAN8 KO cell lines. A representative dot blot is shown in **C**. Each dot represents the signal from an independent sample. Grouped in different colors are secretion assays that were processed in parallel. Total number of replicates are 9. Red dots represent the mean +/- the standard deviation. The p values from a one-way ANOVA analysis with orthogonal contrasts are shown. **E**. Quantification of the number of granules detected in WT (green) and TSPAN8 KO (clone 14) (purple) cells before (2 minutes) and during ATP-stimulation (24 minutes). Representative still frames before and after ATP treatment of the time-lapse are shown in **F**. A representative time-lapse is shown in supplementary video 2. Each dot represents the mean proportion of granules in cells. The gray shadow represents the mean +/- the standard deviation. **F**. Still frames from a time-lapse spinning disk microscopic analysis of mucin secretion in WT (top panels) and TSPAN8 KO (bottom panels) cells. Left and right images are before and after ATP stimulation respectively. Signal is from mucin5AC·GFP fluorescent emission. Intensity was color-coded in and pseudo-color look up table (LUT). Each dot is a mucin5AC·GFP granule (arrowheads). Arrows point to areas of maximal mucin release into the extracellular space. Z-stack images were re-sliced using Fiji/ImageJ to show maximal projection of the lateral view of the imaged field.

To determine how Tspan-8 affected stimulated secretion, we imaged mucin-5AC·GFP in live WT and TSPAN8 KO (clone14) cells by spinning disk microscopy during ATP-stimulated mucin secretion. WT and TSPAN8 KO cells responded similarly during the fast-rate secretory phase by releasing 20 % of mucin-5AC·GFP granules within the first minute after ATP addition (Figure 2E and F; Supplementary Video 2). During the slow-rate of stimulated secretion, WT cells released mucin-5AC·GFP granules at a rate of 0.20 % each minute (linear model: percentage of granules ∼ time; p < 0.001; R^2^: 0.907). After 24 minutes, WT cells had secreted 5 % of the total amount of granules during the second phase of secretion (Figure 2E and F; Supplementary Video 2). TSPAN8 KO cells on the other hand, doubled the rate of release to 0.42 % of granules each minute (linear model: percentage of granules ∼ time; p < 0.001; R^2^: 0.951). After 24 minutes TSPAN8 KO cells had secreted 10 % of the total amount of granules during the second phase of secretion (Figure 2E and F; Supplementary Video 2).

### Tetraspanin-8 is localized at the plasma membrane

Transient transfection of Tspan-8·RFP, immunofluorescence of the plasma membrane-localized sodium/potassium-transporting ATPase (Na+/K+-ATPase) and the analysis of the Pearson Correlation Coefficient (PCC) showed a very high degree of co-localization (Figure 3 A and B). To exclude the possibility that the localization of Tspan-8 could be affected by over expression or transfection, we tagged the c-terminus of the TSPAN8 locus with the sfGFP sequence using CRISPR/Cas9 genome editing (Figure Supplementary 4A). A mucin secretion assay and dot blot analysis showed that the c-terminus tagging of TSPAN8 did not affect mucin production (Figure Supplementary 4B) or secretion (Figure Supplementary 4C and D). Live cell imaging of the Tspan-8 c-terminally tagged with sfGFP co-localized with the lipophilic plasma membrane-marker CellBrite® (Figure 3C; arrows). Immunolabeling of endogenous Tspan-8 in WT cells confirmed the localization at the plasma membrane and absence from mucin granules (Figure Supplementary 4E; arrows). An intracellular pool of Tspan-8 co-localized with the recycling endosomes marker Rab11A (Figure Supplementary 4F; arrowheads) describing the known recycling behaviour of plasma membrane proteins (33, 34).

**Figure 3.**
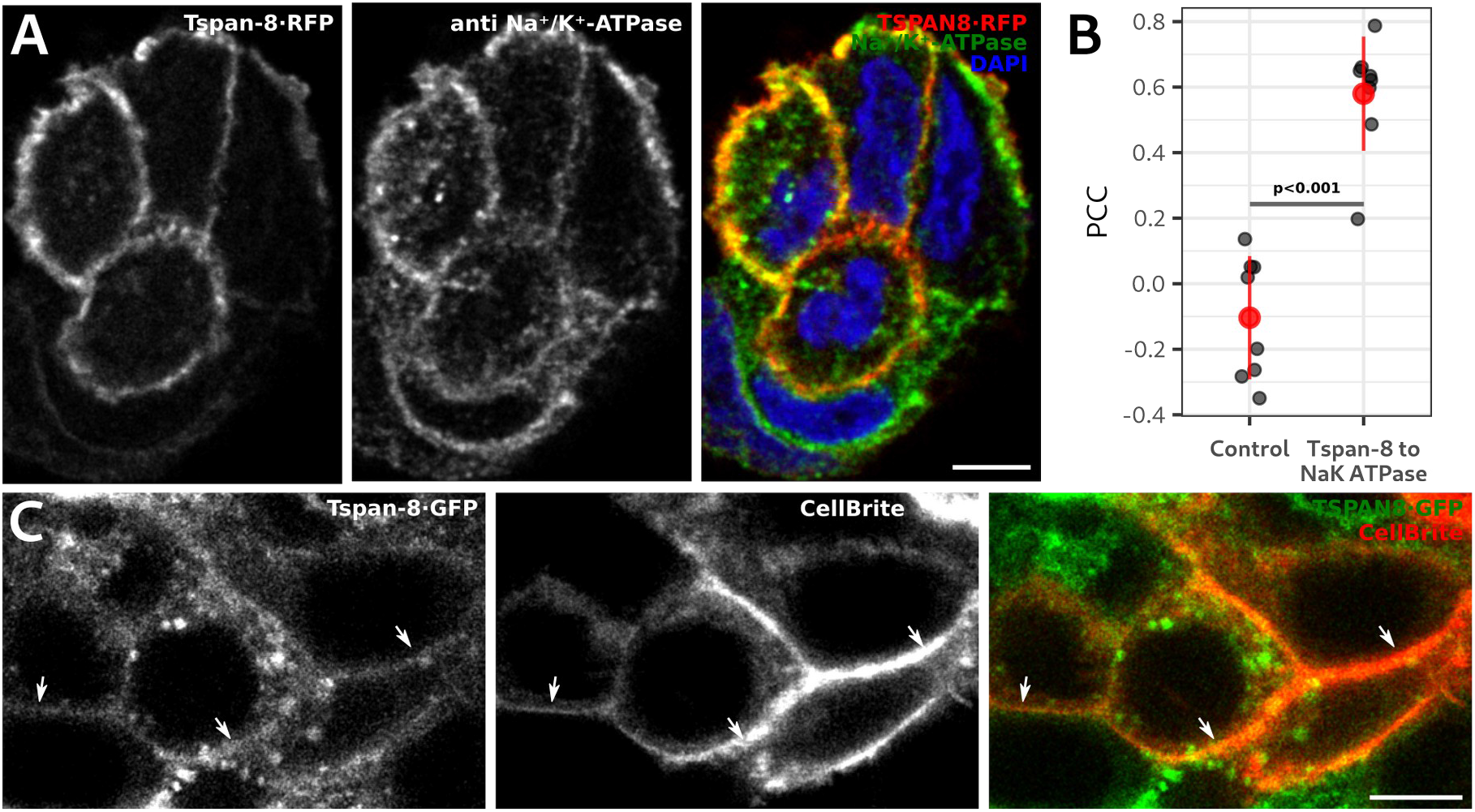
Tetraspanin 8 is localized at the plasma membrane of mucin-secreting cells. **A**. Optical plane of a representative confocal image of mucin-secreting cells transfected with Tspan-8·RFP (left image) and immunolabelled for the Na+/K+-ATPase (middle image). Right image shows the merge of the two and DAPI to visualize the cell nucleus. Scale bar is 10 µm. **B**. Quantification of the Pearson Correlation Coefficient (PCC) between Tspan-8·RFP and Na+/K+-ATPase. A representative image is shown in **A**. As control, Na+/K+-ATPase images were rotated 90º right and the PCC was calculated. Each individual gray dot represents the PCC of one image with at least three cells. Total number of analyzed images are 9. The red dot is the mean value of the gray dots +/- the standard deviation. The p value was calculated with an ANOVA test. **C**. Optical plane from a confocal image of live mucin-secreting cells expressing Tspan-8·GFP at endogenous levels (left image). To visualize the plasma membrane, cells were incubated with the lipophilic membrane marker CellBrite® (middle image). Right image shows the merge of the two. Arrows point to the plasma membrane as determined by the CellBrite® signal. Scale bar is 10 µm.

### Tetraspanin-8 interacts with the Q-SNAREs syntaxin-2 and syntaxin-3

The localization of Tspan-8 and its effect on mucin secretion suggested that it might function by interacting with a plasma membrane-localized syntaxin. We immunoprecipitated Tspan-8·GFP from differentiated mucin-secreting cells and determined by western blot whether syntaxins 1, 2, 3 or 4 were present in the immunoprecipitated samples. Only synatxins 2 and 3 were found to co-immunoprecipitate with Tspan-8·GFP (Figure 4A). We confirmed that Stx2 interacts with VAMP-8 by co-immuno-precipitation of GFP·Stx2 followed by western blot analysis of samples from WT HT29 cells transiently transfected with GFP·Stx2 (Figure 4B). Surprisingly, when we immunoprecipitated Tspan-8·GFP, we detected Stx2 in the protein complex but not VAMP-8 (Figure 4C and Supplementary 5A and B). Munc18-1 and -2 bind to syntaxins to regulate the formation of SNARE complexes (22, 35) and are known for their involvement in mucin secretion (36). By transiently co-transfecting Tspan-8·GFP and Stx2·FLAG and immunoprecipitating GFP, we reconfirmed Stx2 as a Tspan-8 interacting protein, but neither Munc18-1 (Figure 4D) nor Munc18-2 (Figure 4E) were found as part of the Tspan-8/Stx2 protein complex. Stx2 therefore interacts with either VAMP-8 (Figure 4B) or with Tspan-8 (Figure 4C), but not with both concomitantly (Figure 4C and Supplementary 5A and B). To confirm this observation, we transiently co-transfected cells with GFP·Stx2 and Tspan-8·RFP and immunoprecipitated GFP to precipitate the two protein complexes: Stx2/Tspan-8 and Stx2/VAMP-8. We found that VAMP-8 and Tspan-8·RFP both co-immuno-precipitated with GFP·Stx2 (Figure Supplementary 5C).

**Figure 4.**
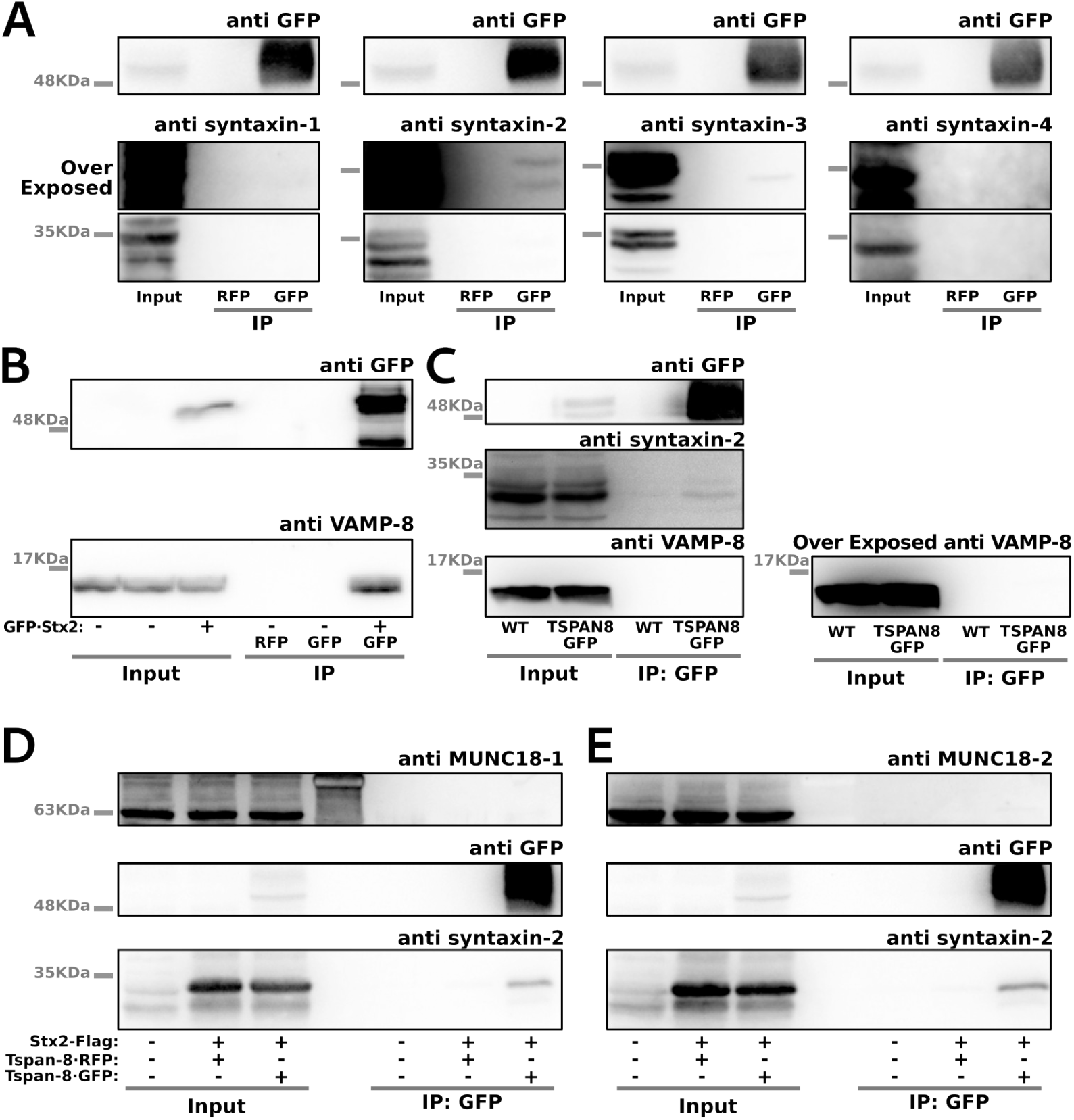
Tspan-8 binds with syntaxins 2 and 3. **A**. Lysates of HT29 cells genetically modified by CRISPR/Cas9 to express Tspan-8·GFP were processed for immunoprecipitation of GFP and western blot analysis. The sample was divided in four different parts. Top panels show immunoblotting against GFP to confirm the immunoprecipitation. Lower panels show the immunoblotting for syntaxins 1 to 4. Middle panels are over-exposed membranes for better visualization of the co-immuno-precipitated proteins. RFP immunoprecipitation was used as control condition. **B**. Lysates of HT29 cells transiently transfected with GFP·Stx2 were processed for immunoprecipitation of GFP and western blot analysis. Top panel shows immunoblotting against GFP to confirm the immunoprecipitation. Lower panel shows the immunoblotting against VAMP-8. Untransfected cells and RFP immunoprecipitation were used as control conditions. **C**. Lysates of HT29 cells genetically modified by CRISPR/Cas9 to express Tspan-8·GFP were processed for immunoprecipitation of GFP and western blot analysis. Top-left panel shows immunoblotting against GFP to confirm the immunoprecipitation. Middle-left panel shows the immunoblotting against Stx2. Bottom-left panel shows immunoblotting against VAMP-8. Bottom-right panel is the over-exposed VAMP-8-immunoblotted membrane. WT HT29 (no GFP) cells were used as a control condition. **D – E**. Lysates of WT HT29 cells transiently co-transfected with Tspan-8·GFP and Stx2·FLAG were processed for GFP immunoprecipitation and western blot analysis. Top panels show immunoblotting against Munc18-1 (**D**) and Munc18-2 (**E**). Middle panels show immunoblotting against GFP to confirm the immunoprecipitation. Lower panels show the immunoblotting against Stx2. No transfection and Tspan8·RFP transfection were used as control conditions.

### Syntaxin-2 is the plasma membrane Q-SNARE required for mucin secretion

Immunolabelling of Stx2 and the Na+/K+-ATPase in HT29/Caco-2 co-cultures and quantification of the PCC in each optical slide, showed that Stx2 localized to the apical region of the plasma membrane (Figure 5A and B). TSPAN8 KO did not affect the localization of Stx2 (Figure Supplementary 6A and B). The PCC between Stx2 and mucin-5AC·GFP showed that Stx2 does not localize to mucin granules (Figure 5B and Supplementary 6A – C). STX2 gene ablation (Figure Supplementary 6D) did not affect mucin-5AC levels (Figure 6C). A secretion assay showed that STX2 removal reduced the ATP-stimulated secreted mucin-5AC·GFP by 64% (Figure 5D and E). STX2 RNAi treatment in WT HT29 cells confirmed this finding (Figure 6C and D). Basal mucin secretion was also decreased in the STX2 KO (Figure 5D and E). Quantification of beta-tubulin in the cell lysates by western blot shows equal amount of cells in these secretion assays (Figure Supplementary 6E and F). These data confirm the involvement of Stx2 in mucin secretion. The residual amount of mucin secretion in the STX2 KO cells suggests some level of redundancy, and it is likely that syntaxin-3 also functions in mucin secretion. This fits well with the recent findings of Brunger and colleagues where they use syntaxin-3, SNAP-23 and VAMP8 in an *in vitro* reconstituted, membrane fusion assay to recapitulate aspects of mucin granule fusion (37).

**Figure 5.**
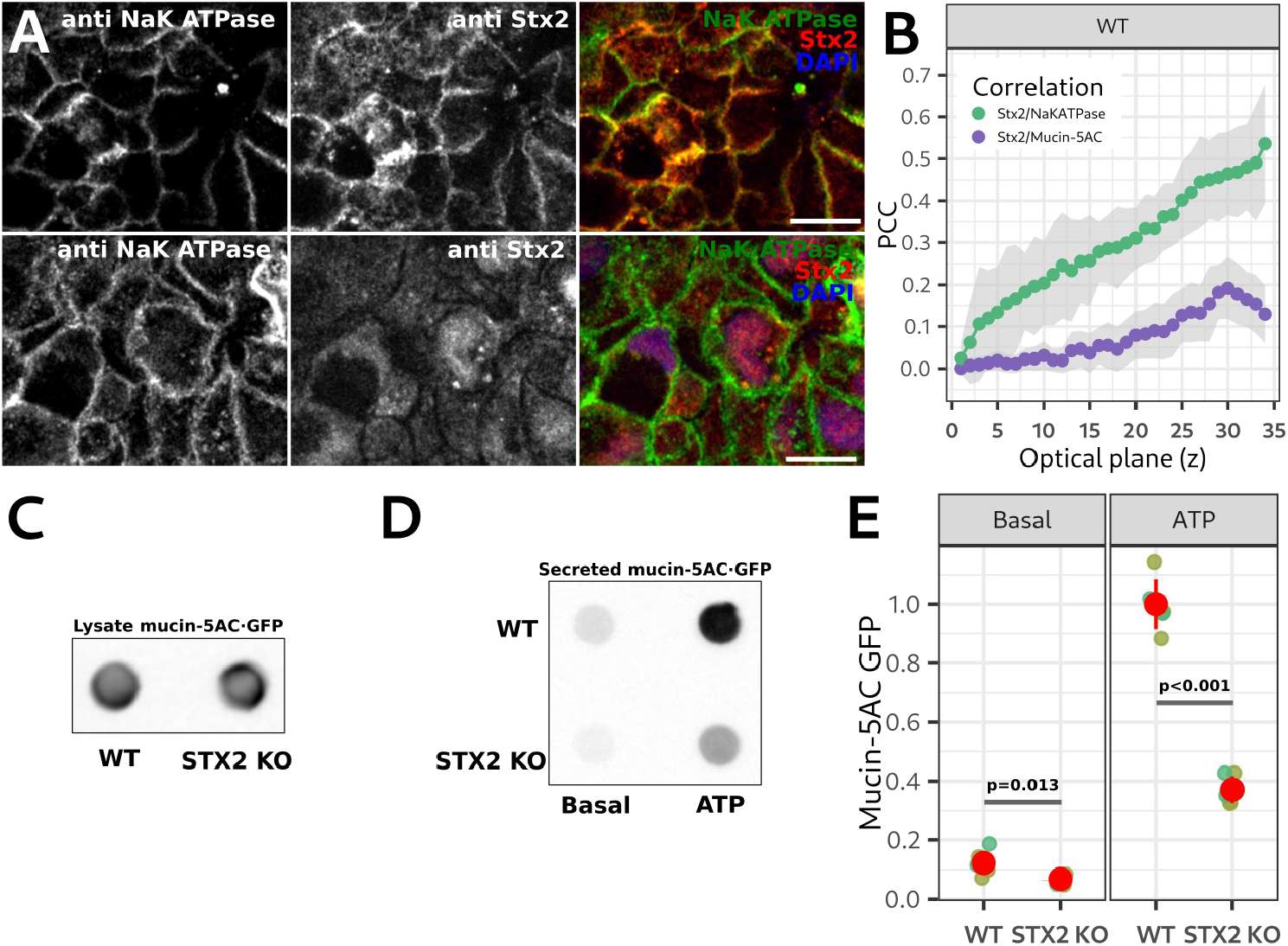
Syntaxin-2 in necessary for mucin-5AC secretion. **A**. Optical planes of a representative confocal image of a HT29 WT/Caco2 co-culture immunolabelled for the Na+/K+-ATPase (left images) and Stx2 (middle images). Right images show the merge of the two and DAPI to visualize the cell nucleus. Scale bar is 10 µm. Top row is an optical plane of the apical part of the co-culture and the lower row is an optical plane of the basal/medial part of the co-culture and distinguishable by the presence of DAPI staining. HT29 (clone 10) cells express mucin-5AC·GFP (shown in Figure supplementary **6C**) at endogenous levels. **B**. Green dots and lines show the mean PCC quantification between the Na+/K+-ATPase and Stx2 signals in each optical plane of HT29 WT/Caco2 co-cultures. Representative images are shown in **A**. Purple dots and lines show the mean PCC between Stx2 and mucin-5AC·GFP. A representative image is shown in Figure Supplementary **6C**. The gray areas show the mean PCC +/- the standard deviation. **C**. Representative dot blot showing the total (cell lysate) mucin-5AC·GFP content in differentiated WT and STX2 KO cell lines. Mucin-5AC.GFP signal was detected from sfGFP fluorescence emission. Left dot is a sample from WT cells and the right dot is a sample from the STX2 KO cell line. **D**. Representative dot blot of a secretion assay of WT and STX2 KO cells. sfGFP fluorescence emission is shown. **E**. Quantification of secretion assays of non-stimulated (basal) or ATP-stimulated WT and STX2 KO cell lines. A representative dot blot is shown in **D**. Each dot represents the mucin-5AC·GFP signal from a single secretion assay. Grouped in different colors are secretion assays that were processed in parallel. Total number of replicates are 9. Red dots represent the mean +/- the standard deviation. Values are expressed as relative to the average mucin-5AC·GFP signal in ATP-stimulated WT cells. Statistical analysis was performed independently for basal and ATP-stimulated conditions. The p values from a one-way ANOVA analysis with orthogonal contrasts are shown.

**Figure 6.**
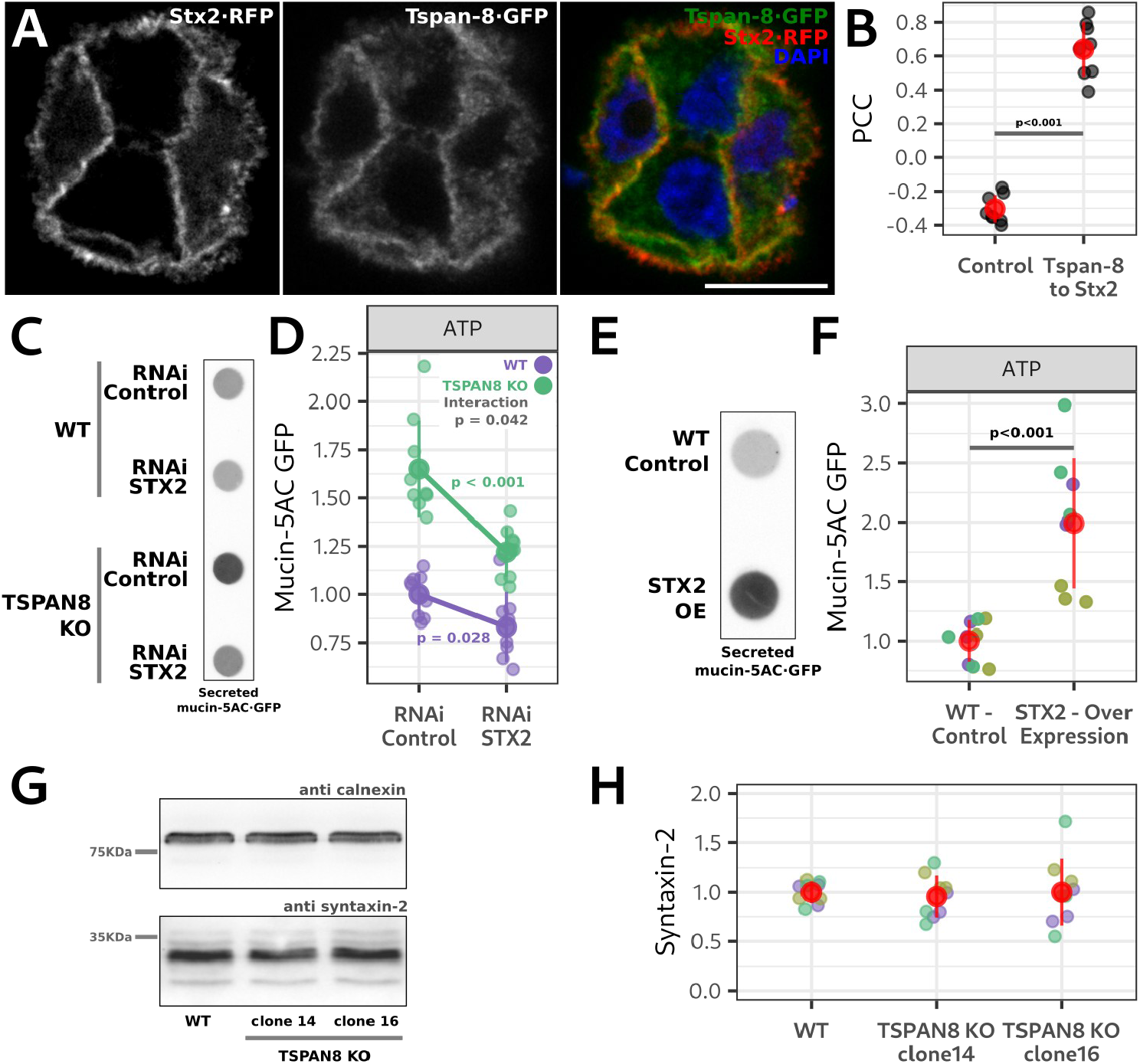
Syntaxin-2 and Tspan-8 are in the same pathway of regulated mucin secretion. **A**. Optical plane of a representative confocal image of cells co-transfected with Stx2·RFP (left image) and Tspan-8·GFP (middle image). Right image shows the merge of the two and DAPI to visualize the cell nucleus. Scale bar is 10 µm. **B**. Quantification of the PCC between Stx2·RFP and Tspan-8·GFP. A representative image is shown in **A**. As control, Stx2·RFP images were rotated 90º right and the PCC was calculated. Each individual gray dot represents the PCC of one image with at least three cells. Eight independent images were analyzed. The red dot is the mean value of the gray dots +/- the standard deviation. The p value of an ANOVA test is shown. **C**. Representative dot blot showing ATP-stimulated mucin-5AC·GFP secretion from WT and TSPAN8 KO cells treated with RNAi control and RNAi against STX2. **D**. Quantification of the amount of mucin-5AC·GFP from ATP-stimulated WT and TSPAN8 KO cells treated with RNAi control and RNAi against STX2. A representative dot blot is shown in **C**. Each dot represents the signal from a single secretion assay. Total number of replicates are 9. Bigger solid-filled dots represent the mean +/- the standard deviation. A two-way ANOVA with interaction was done. Genotype:Treatment interaction p value = 0.042; Genotype principal factor p value < 0.001; Treatment principal factor p value < 0.001. **E**. Representative dot blot of a secretion assay showing the secreted mucin-5AC·GFP of ATP-stimulated WT and STX2-over expressing (OE) cells. **F**. Quantification of the secretion assays of ATP-stimulated WT and STX2-over expressing cells. A representative dot blot is shown in **E**. Each dot represents the mucin-5AC·GFP signal from a single secretion assay. Grouped in different colors are secretion assays that were processed in parallel. Total number of replicates are 9. Red dots represent the mean +/- the standard deviation. A one-way ANOVA was done and the p value is shown in the graph. **G**. Representative western blot of the total amount (lysate) of Stx2 (lower panel) and calnexin (upper panel) in WT and TSPAN8 KO cells. Membranes were immunoblotted with anti calnexin and anti Stx2 antibodies, respectively and developed by ECL. **H**. Quantification of the amount of Stx2 in WT and TSPAN8 KO cells. A representative western blot is shown in **G**. Each dot represents the Stx2 signal from an independent sample. Grouped in different colors are samples that were processed in parallel. Total number of replicates are 9. Red dots represent the mean +/- the standard deviation. A one-way ANOVA analysis was done, and no statistical differences were found. Genotype principal factor p value = 0.913.

### Syntaxin-2 and Tspan-8 are in the same pathway of regulated mucin secretion

Tspan-8 and Stx2 both localized to the plasma membrane (Figure 3 and Figure 5) and co-immuno-precipitated (Figure 4). Co-transfection of Stx2·RFP and Tspan-8·GFP and quantification of the PCC between them confirms they are co-localized together at the plasma membrane (Figure 6A and B).

STX2 RNAi in WT and TSPAN8 KO cells decreased STX2 expression between 50 to 60% on average (Figure Supplementary 7A and B). Acute downregulation of STX2 by RNAi did not affect mucin-5AC·GFP expression (Figure Supplementary 7C – F). We found that STX2 downregulation differentially decreased mucin-5AC·GFP release in WT and TSPAN8 KO cells (Figure 6C and D – ANOVA interaction p value = 0.042; Loading controls in Figure Supplementary 7G and H) suggesting that both proteins function in the same pathway leading to mucin secretion.

To provide further functional evidence that the amount of Stx2 available at the plasma membrane regulates the quantity of mucin release, we over expressed STX2 with a lentiviral system in mucin-secreting cells. Puromycin-selected and differentiated cells had an average 3-fold increase in STX2 expression (Figure Supplementary 7I and J). STX2 over expression did not change mucin-5AC·GFP expression (Figure Supplementary 7K – N) but doubled the amount of mucin-5AC·GFP secreted in ATP-stimulated cells (Figure 6E and F; Loading controls in Figure Supplementary 7O and P).

Since over expression of STX2 increases mucin release we determined by western blot whether TSPAN8 KO affected STX2 expression. We found that Tspan-8 ablation did not change STX2 levels (Figure 6G and H).

## Discussion

An understanding of the mechanisms by which cells control the release propensity, particularly in the two phases of stimulated secretion, is an important unaddressed issue. Our data show that TSPAN8 expression is up-regulated in mucin-secreting goblet cells and that Tspan-8 functions at the plasma membrane to control the release propensity of mucin granules during the slow and sustained phase of ATP-stimulated secretion. Identification of tetraspanin-8 in controlling mucin release by sequestering syntaxin-2 answers the long-standing question of how cells regulate the external agonist-stimulated, biphasic release of secretory cargoes.

### Tspan-8 controls the number of mucin granules that can dock for fusion

External agonist-stimulated fusion of secretory granules to the plasma membrane is biphasic. The first high-rate secretion phase lasts a few seconds and the second phase has a lower release rate but can last several minutes (14, 38). Docking is a fundamental process preceding vesicle fusion that is mediated by syntaxins (26-28). It has been suggested that docking of synaptic vesicles during sustained neurotransmitter release is the rate limiting step (17).

Our data reveal that stimulated mucin release is biphasic and that TSPAN8 is crucial to control the release propensity of granules in reserve. TSPAN8 KO increased mucin release during the second phase of secretion (Figure 2). Syntaxin-2 localization and expression are not affected by TSPAN8 KO (Figures 5 and 6), therefore Tspan-8 controls the quantities of syntaxin-2 available at the plasma membrane for docking of granules (Figure 7). Tspan-8 localizes to the plasma membrane of secretory cells (Figure 3), co-localizes with Stx2 (Figure 6), and co-immunoprecipitates with syntaxins 2 and 3, but not 1 and 4 (Figure 4). Furthermore, we show that Stx2 can form a complex with either VAMP-8 or Tspan-8, but these two complexes are mutually exclusive. The Tspan-8/Stx2 complex does not contain VAMP-8 or Munc18 (Figure 4). This strongly suggests that ordinarily, Tspan-8 regulates secretion by binding or sequestering Stx2 (and perhaps Stx3) to effectively limit the available Q-SNAREs at the plasma membrane and keeping docking sites empty and refractory (not accessible for an incoming vesicle). Cells lacking Tspan-8 provide quantitatively more Stx2 for docking and therefore increases the fusion of mucin granules. The kinetics of the fast-rate phase of mucin release in WT and TSPAN8 KO cells are similar (Figure 2). Tspan-8 therefore does not affect mucin granules that are docked before ATP stimulation but prevents the docking of a pool of granules that are kept in reserve (Figure 7). This also explains why Tspan-8 loss does not affect baseline mucin secretion.

**Figure 7.**
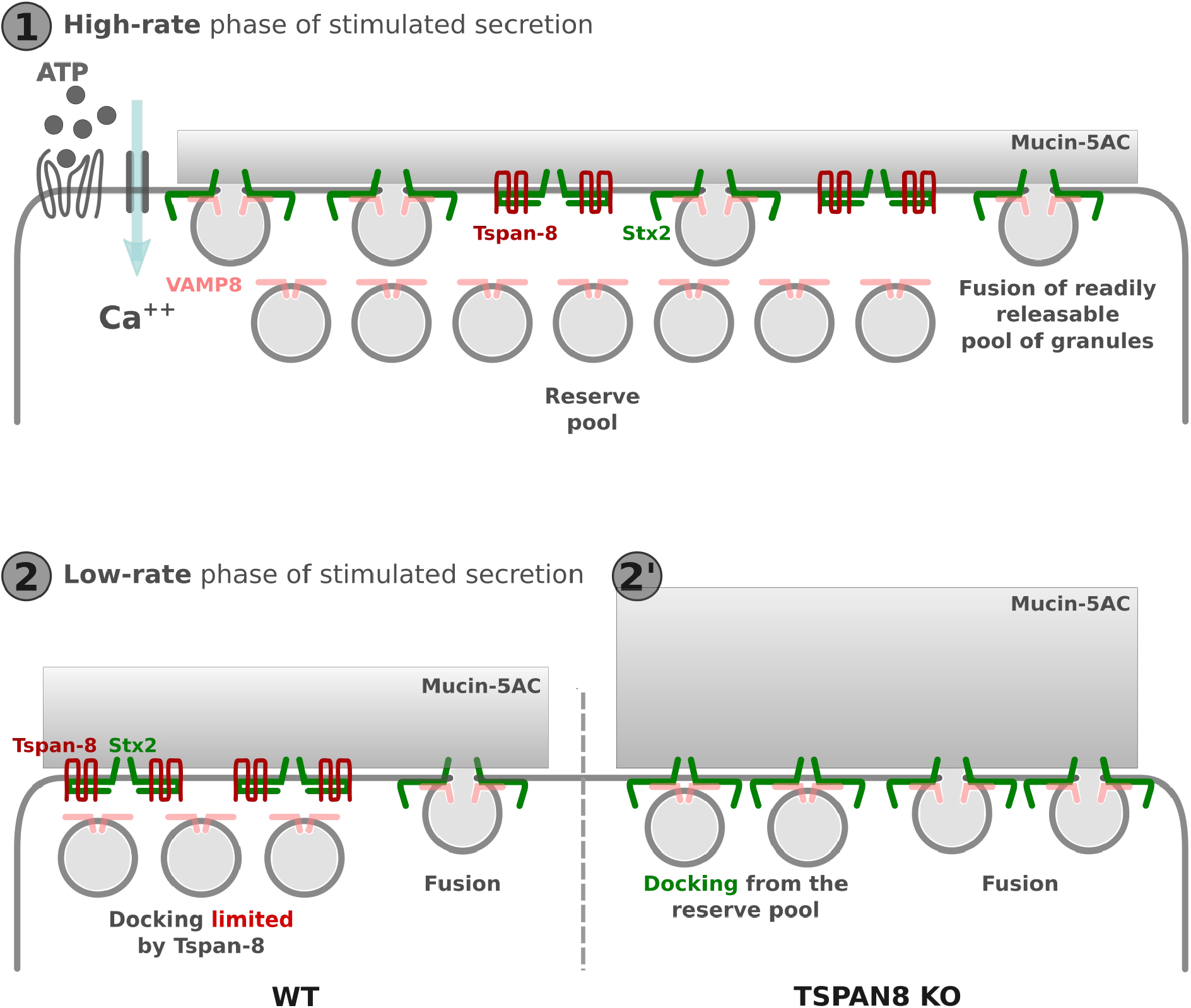
Model for function of Tspan-8 to regulate biphasic release of granules. 1. Pre-docked granules fuse upon exposure of cells to external agonist like ATP. **2**. Tspan-8 at the plasma membrane binds Stx2 preventing it from docking granules from the reserve pool during the second phase of stimulated secretion. **2’**. Loss of Tspan-8 exposes Stx2 to engage with granules from the reserve pool that fuse to increase the quantity of mucins secreted.

### A general mechanism for docking granules in reserved pool

It is tempting to propose that Tspan-8 or a different tetraspanin modulates other granule secretion, including insulin or neurotransmitters in a similar manner. Insulin secretion is also biphasic (38). Tspan-8 is expressed in the human pancreas (39) and allelic variants of Tspan-8 have been associated with type-II diabetes mellitus (40, 41). Future studies should determine if Tspan-8 is involved in insulin secretion in human beta cells and its impact in type-II diabetes. Mouse beta cells on the other hand, do not express Tspan-8 and a KO model showed no defect in insulin secretion (39). It is conceivable that a different tetraspanin could play a similar role in mice. Tspan-8 is also expressed in the human brain (25), perhaps to modulate sustained neurotransmitter release.

Identification of Tspan-8 in sequestering a Q-SNARE to control biphasic granule release, paves the way to explain and to further understand how cells keep granule in reserve to handle quantities of cargoes secreted based on the cellular needs.

## Methods

### Lead Contact

Requests for further information and or reagents may be addressed to the Lead Contact, Malhotra Vivek.

### Data and Code Availability

The R code to reproduce the bioinformatic analysis of the data published by (29) is available at the publisher website and at https://github.com/boina/wojnacki_etal_2022/

### Cell culture and Differentiation

HT29-N2 cells were grown in Dulbecco’s Modified Eagle’s Medium (DMEM) cell culture media (Lonza. Cat. Nº: BE12-604F/U1) supplemented with 10% (vol / vol) heat-inactivated fetal bovine serum (Gibco. Cat. Nº: 10270-106) (complete media). Cells were passaged at a 1:4 ratio every Monday and Friday. Cells were kept at 37ºC in a humidified incubator supplied with CO2 5%.

For differentiation, 2×10^5^ cells / cm^2^ were plated into a T25 or T75 culture flask with complete media, typically on a Monday. The following day, cells were washed with phosphate buffered saline (PBS) and complete media was changed with Protein-Free Hybridoma Media (PFHM) (Gibco. Cat. Nº: 12040-051). After 3 days, media was replaced with fresh PFHM. After 6 days in culture, differentiated cells were trypsinized (Gibco. Cat. Nº: 25300-054). For secretion assays and live-imaging experiments, 1.1×10^5^ cells / cm^2^ were plated in complete media in 6-or 12-well plates. For live imaging, cells were seeded in glass-bottom 35 mm dishes (ibidi. Cat. Nº: 81156). For immuno-fluorescence, cells were plated at 1.7×10^4^ cells / cm^2^ in culture dishes containing cover-slips for microscopy. For immunoprecipitation experiments 2×10^5^ cells / cm^2^ were plated in 60 mm dishes. The following day, cells were washed with PBS and complete media was changed with PFHM (30, 31).

### HT29-N2 / Caco-2 co-culture

Caco2 and HT29 co-culture were prepared as previously described (Kleiveland C.R.,2015), with small variations. Caco2 cells were cultured in DMEM 4.5 g/L Glucose with UltraGlutamine (Lonza: H3BE12-604F/U1) medium containing 20% fetal calf serum (Gibco: 10270106), supplemented with 100 units/mL penicillin, and 100 μg/mL streptomycin (Invitrogen: 15070063). Cells were incubated at 37 °C in a humidified incubator supplied with 5% CO2. Caco-2 and HT29 cells were mixed prior to seeding in a ratio of 50:1 (Caco-2/HT29) at a density of 2.5 10^5^/cm^2^. The co-culture was maintained in DMEM 4.5 g/L Glucose with UltraGlutamine medium supplemented with 20% fetal calf serum, and the culture medium was changed every 2 – 3 days. Images of polarized HT29 clusters were taken between 14 and 21 days after seeding.

### Cell Transfection

Cells were transfected for transient plasmid over expression with lipofectamine (ThermoFisher Scientific. Cat. Nº: L3000001) as previously described (42) at the moment of plating. For immuno-fluorescence experiments, cells were transfected in 35 mm in diameter cultures dishes with 2 ml of complete culture media. For each transfection reaction 7 µl of lipofectamine were diluted in 125 µl of OptiMEM and between 1.5 to 3 µg of plasmid DNA were diluted in 125 µl of OptiMEM. The lipofectamine and DNA solution were then mixed incubated at RT for 15 minutes and added to the culture dish. For immuno-precipitation transfections the ratio of lipofectamine to DNA were maintained but scaled up to 250 µl for each reaction transfection.

### RNA Interference Transfection

Cells were transfected for RNA interference with lipofectamine RNAiMAX (ThermoFisher Scientific. Cat. Nº: 13778075). HT29 cells were differentiated and seeded in 24-well plates. At the moment of seeding the first RNAi transfection mix was added to the culture media. For each reaction, the transfection mix was prepared by diluting 3.125 µl of lipofectamine in 50 µl of OptiMEM. The RNAi mix was prepared by diluting two independent RNAi against STX2 down to 20 nM each one. The lipofectamine and the RNAi solutions were mixed and incubated at RT for 10 minutes. The mix was then added to the culture well. Two days after the first reaction, a second, identical transfection was performed and 4 days after the first transfection, the secretion assay were performed.

### Sequences of the RNA interference used

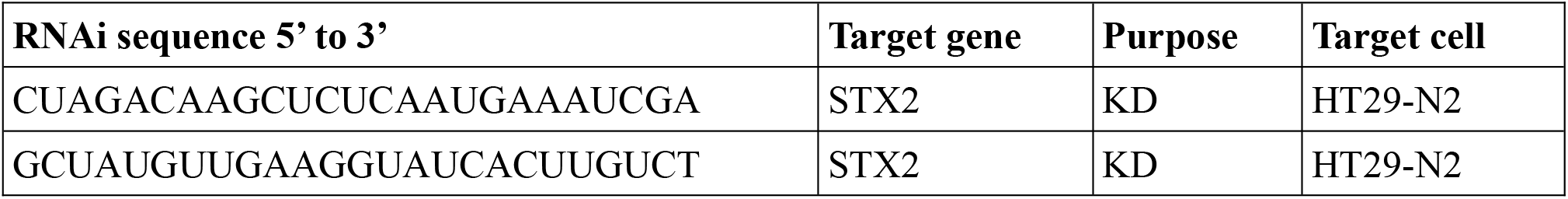

### CRISPR/Cas9 genetic engineering

To generate Knock-Out (KO) and Knock-In (KI) cell lines we used the RNA-guided Cas9 endonuclease system developed from the microbial clustered regularly interspaced short palindromic repeats (CRISPR) adaptive immune system (43). Guide RNA sequences were chosen with the web-based tool CRISPOR which implements scoring algorithms based on their potential off target and on-target DNA cleavage activity (44). The highest-scored guide RNA sequences were cloned into a pSpCas9(BB)-2A-GFP backbone plasmid. pSpCas9(BB)-2A-GFP (PX458) was a gift from Feng Zhang (Addgene plasmid #48138; http://addgene.org/48138; RRID: Addgene_48138) (43). After confirmation of correct insertion of the guide RNA oligonucleotides by colony PCR and sequencing, plasmids were amplified, purified with a midi prep kit and transfected to HT29-N2 cells with lipofectamine 3000 as described in the cell transfection section of materials and methods. Two to 3 days after transfection, cells were trypsinized and detached from the culture dish, pelleted and re-suspended in complete culture media. Cells were then sorted by a Fluorescence Activated Cell Sorter (FACS). For KO generation 1 GFP-positive cell was placed in each wheel of a 96 multi-well plate containing complete media supplemented with plasmocin (15 µg / ml) (InvivoGen. Cat. Nº: ant-mpt). Cells were amplified until final confirmation of gene knockout by RT-PCR and protein electrophoresis followed by western blot as previously described (45). To increase the chances of successful gene KO 3 different guide RNAs targeting the ATG origin of translation, middle and c-terminus regions of the target locus were co-transfected. For gene KI, 1 guide RNA (designed to target the c-terminus of the locus) was co-transfected with the PCR product of the sfGFP sequence amplified with primers bearing tails complementary to the upstream and downstream regions of the expected cut site by Cas9 in the target gene. Two to 3 days after transfection, GFP-positive cells were bulk-collected by FACS and grown for 10 to 15 days in complete media. After this time, GFP signal from the CRISPR/Cas9 plasmid has diluted away and any GFP signal should come from the integration of the transfected DNA template into the target genome. Cells were then sorted again by FACS and 1 GFP-positive cell was placed in each wheel of a 96 multi-well plate containing complete media supplemented with plasmocin (15 µg / ml). Cells were amplified until final confirmation of gene KI by PCR and protein electrophoresis followed by western blot.

### List of guide RNA sequences

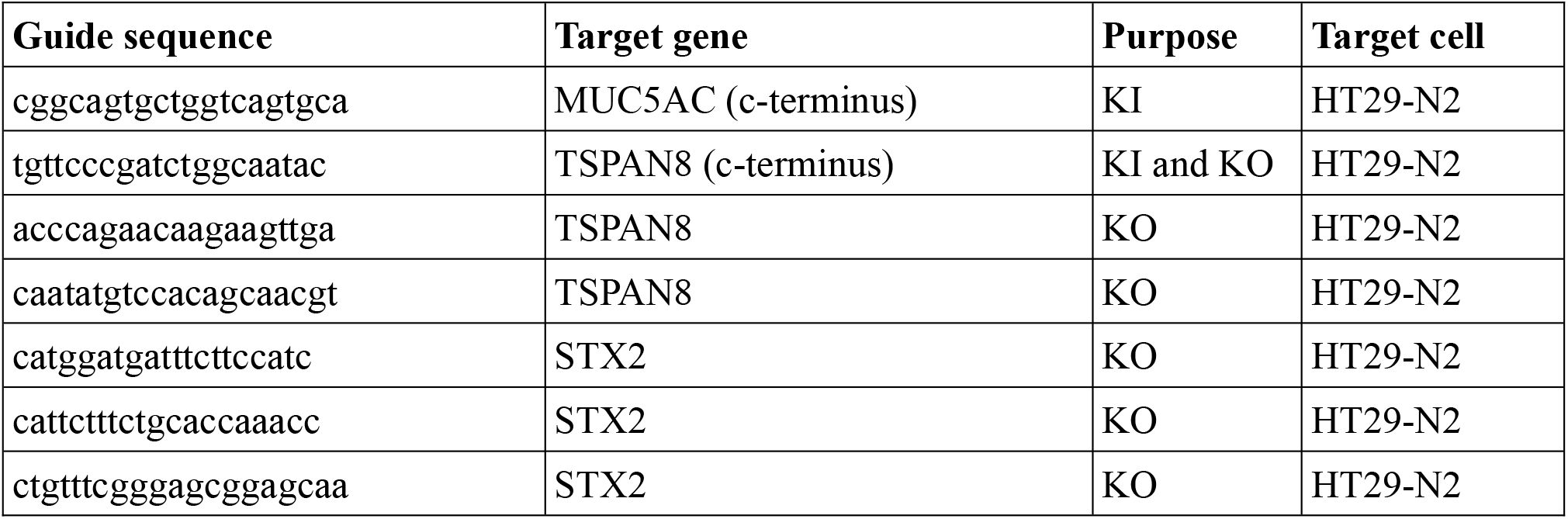

### Generation of stable over expressing cell lines

Lentiviral particles were generated by co-trasnfecting pRSV–REV, pMDLg/pRRE, VSV-G (46) and the transfer plasmid (L309) carrying the sequence of the gene to be over expressed and resistance to puromycin into 0.8×10^6^ HEK293 cells in a 60 mm culture dish with 6 µl of TrasnIT (Cat. Nº: Mirus Bio. 293) containing 3 ml of complete media. The following day the cell culture media was changed with fresh media. Two and three days after, the cell media was colected and filtered through a 0.45 µm membrane. HT29 cells were plated in 60 mm dishes. The following day the media was changed for 2 ml of fresh complete media, 1 ml of the filtered viral particles and 8 µg/ml of hexadimethrine bromide (polybrene) (Cat. Nº: Sigma-Aldrich. 107689). The following day media was change with fresh media. Two days after, the culture media was changed for complete media supplemented with puromycin (Cat. Nº: Gibco. A11138-03) 15 µg/ml for 1 week. After this time, cells were maintained in compete media supplemented with puromycin 2.5 µg/ml.

### Secretion Assays

Secretion assays were done as previously described (3, 31, 47, 48). Culture media of differentiated HT29-N2 cells was replaced with 0.5 ml (12-well plates) or 1 ml (6-well plates) of isotonic buffer (KCl 2.5mM; NaCl 140 mM; CaCl_2_ 1.2 mM; MgCl_2_ 0.5 mM; Glucose 5 mM; HEPES 10 mM; pH 7.42 adjusted with tris base and 305 mOsm / litre adjusted with D-manitol if needed). For stimulated secretion, the isotonic solution was supplemented with 100 µM ATP. Cells were kept with the isotonic solution at 37ºC for 30 minutes. The supernatant was collected and centrifuged for 5 minutes at 800 g at 4ºC. 80 % of the supernatant was recovered in a new tube (secreted fraction). Cells were lysed in lysis buffer as described in the western blot section of material and methods.

### Dot blot

100 microlitres of the secreted fractions and 100 µl of 1:20 diluted (in PBS) lysates from the secretion assays were loaded into a bio-blot microfiltration blotting system (BioRad. Cat. Nº: 1706545). Samples were left to flow through a 0.45 µm nitrocellulose blotting membrane (Amersham Protran. Cat. Nº: 10600002) by gravity (3, 47). Membranes were then blocked with a Tris-Buffered Saline (TBS) solution supplemented with 0.1 % vol / vol polysorbate 20 (Sigma-Aldrich. Cat. Nº:P1379) (TBST) and 2.5 % weight / vol non-fat dry milk for 20 – 40 minutes at room temperature (RT) on a laboratory rocker. Primary antibodies were diluted in TBST / BSA 5 % (weight / vol) and membranes were incubated with this solution overnight at 4ºC on a laboratory rocker. Fluorescent secondary antibodies (donkey anti rabbit IgG – Alexa Fluor 680 – Life technologies. Cat. Nº: A10043 or donkey anti mouse IgG – Alexa Fluor Plus 800 – Invitrogen. Cat. Nº: A32789) were diluted to 2 µg / µl in TBST / non-fat dry milk 2.5 % and membranes were incubated with this solution for 60 minutes at RT on a laboratory rocker. Fluorescent signal was detected using the iBright imaging system (ThermoFisher Scientific) equipped with a high resolution (9.1 mega pixels) CCD camera and multiplexed laser excitation / emission filters.

### Immunoprecipitation

Cells were washed once with cold PBS and 1.2 ml of lysis buffer (Tris 50 mM; NaCl 150 mM; pH 7.3; Triton X-100 1 % (Sigma-Aldrich. Cat. Nº: T9284); Leupeptin 5 µM (Focus Biomolecules. Cat. Nº: 10-1346); Aprotinin 2 µg/µl (Abcam. Cat. Nº: ab146286); Pepstatin A 2 µg/µl (Panreac Química. Cat. Nº: A2205)) was added. Cells were kept in the lysis buffer on ice for 10 minutes and on a laboratory rocker. Cells were then flushed by gentle pipetting, collected in 1.5 ml tubes and placed in a rotating wheel at 4ºC for 30 minutes. Samples were centrifuged at 12000 g at 4ºC for 15 minutes. 0.1 ml of the supernatant was recovered in a new tube and 20 µl of 6X loading buffer (Tris – HCl 375 mM; SDS 12 %; Glycerol 60 %; DTT 600 mM; Bromophenol Blue 0.6 %) was added (input). One ml of the supernatant was recovered in a new tube containing 25 µl of sepharose beads covalently attached to anti GFP nanobodies (VHH) (ChromoTek. Cat. Nº: gta). Samples were incubated in a rotating wheel for 60 minutes at 4ºC. Beads were precipitated by centrifugation at 2500 g for 5 minutes and washed with 500 µl lysis buffer 4 times. Beads were resuspendended in 30 µl 1.2X loading buffer (IP fraction).

### Protein electrophoresis and western blot

Cells were washed once with cold PBS and 0.5 ml (12-well plates) or 1 ml (6-well plates) of lysis buffer (Tris 50 mM; NaCl 150 mM; pH 7.3; Triton X-100 1 % (Sigma-Aldrich. Cat. Nº: T9284); Leupeptin 5 µM (Focus Biomolecules. Cat. Nº: 10-1346); Aprotinin 2 µg/µl (Abcam. Cat. Nº: ab146286); Pepstatin A 2 µg/µl (Panreac Química. Cat. Nº: A2205); DTT 1 µM (Sigma-Aldrich. Cat. Nº: 43815)) was added. Cells were kept in the lysis buffer on ice for 5 minutes and on a laboratory rocker. Cells were then flushed by vigorous pipetting (12-well plates) or scraped (6-well plates or 60 mm dishes), collected in 1.5 ml tubes and placed in a rotating wheel at 4ºC for 20 – 30 minutes. Samples were centrifuged at 12000 g at 4ºC for 15 minutes. The supernatant was recovered in a new tube and 1:6 in volume of 6X loading buffer was added (lysate). Lysates with loading buffer were boiled at 95ºC for 5 minutes. Twenty to 40 µl of the lysate were loaded into a 12 % polyacrylamide gel for SDS-PAGE electrophoretic (BioRad. Mini-PROTEAN) separation of proteins. Samples were run 20 minutes at constant 90 V then 110 V for 60 to 80 minutes. Proteins were transferred to a 0.45 µm PVDF membrane (Amersham. Cat. Nº: 10600023) for 90 minutes at constant 400 mA in an ice-bucket. Membranes were blocked with a solution of 2.5 % (weight / weight) of non-fat dry milk in TBST for 20 to 30 minutes at RT on a laboratory rocker. Primary antibodies were diluted in TBST supplemented with 5 % BSA and membranes were incubated with this solution overnight at 4ºC. Membranes were washed 3 times, 5 minutes each. Horseradish root peroxidase-conjugated (Jackson ImmunoResearch) or fluorescently-conjugated (Invitrogen) secondary antibodies were diluted in TBST to 0.08 µg/µl and 0.2 µg/µl respectively. Membranes were incubated with the secondary antibody solution for 60 minutes at RT in a laboratory rocker. Membranes were washed 3 times, 5 minutes each with TBST. Enhanced chemiluminescence (ECL) signal was generated by incubating the membranes with Immobilon Forte Western HRP Substrate (Millipore. Cat. Nº: WBLUF0100). ECL and fluorescent emission were digitalized with the iBright imaging system (ThermoFisher Scientific) equipped with a high resolution (9.1 mega pixels) CCD camera.

### Immunofluorescence

Cells were fixed for 15 minutes with a solution of paraformaldehyde 4 % (weight / vol) (Sigma-Aldrich. Cat. Nº: P6148) in PBS, pre-warmed at 37ºC. Cells were then washed with PBS 3 times, 5 minutes each and non-specific binding sites were blocked with a solution of saponin (Sigma-Aldrich. Cat. Nº: 47036) 0.05 % (weight / vol) / bovine serum albumin (Sigma-Aldrich. Cat. Nº: A7906) 0.2 % (weight / vol) in PBS (blocking solution from now on) for 20 minutes at RT. Primary antibodies were diluted in blocking solution and cells were incubated with this solution over-night at 4ºC. After primary antibody incubation, cells were washed 3 times with blocking solution, 5 minutes each. All secondary antibodies were diluted to 2 µg/µl in blocking solution. Cells were incubated with the secondary antibodies’ solution for 1 hour at RT. Cells’ nuclei were stained with 4’, 6-diamidino-2-phenylindole, dihydro-chloride (DAPI) (Invitrogen. Cat. Nº: D3571) at 0.5 µg/µl. DAPI was added to the secondary antibodies’ solution. Finally, cells were washed 3 times with blocking solution, 3 times with PBS, rinsed in Milli-Q water once, and mounted on a microscopy slide with FluorSave® reagent (CalBiochem. Cat. Nº: 345789). When antibody-independent labelling of the plasma membrane was required, after the incubation with the secondary antibodies, cells were washed 3 times, 5 minutes each with PBS and incubated for 10 minutes at RT with a 1:200 dilution in PBS of the lipophilic dye CellBrite™ (Biotium. Cat. Nº: 30023). Finally, cells were washed 3 times with PBS and mounted on a microscopy slide with FluorSave.

### List of antibodies and dyes

When the concentration of the stock antibody or the dye solution were not available, the dilution is declared.

| **Antibody or dye** | **Manufacturer** | **Reference** | **Western or dot blot concentration (µg / µl)** | **IF concentration (µg / µl)** |
| --- | --- | --- | --- | --- |
| Anti Munc18-2 | Abcam | ab103976 | 1 | NA |
| Anti Sodium Potassium ATPase | Abcam | ab283318 | NA | 5 |
| Anti syntaxin-3 | Abcam | ab133750 | NA | 10 |
| Anti Tspan-8 | Abcam | ab70007 | 1 | 10 |
| Anti VAMP-8 | Abcam | ab76021 | 0.2 | NA |
| Anti TGN46 | abd serotec | AHP500 | NA | 1 |
| CellBrite Red | Biotium | BT-30023 | NA | 1:200 |
| Anti Rab11A | Cell Signaling | 2413 | 0.5 | NA |
| Anti syntaxin 1 | Chemicon | AB5820 | 1:1000 | NA |
| Anti syntaxin-3 | Chemicon | AB5448 | 1:1000 | NA |
| Anti syntaxin 4 | Chemicon | AB5330 | 1:1000 | NA |
| Anti RFP | Evrogen | AB233 | 20 | NA |
| Anti mucin5-AC | GeneTex Inc | GTX75418 | 0.04 | 0.2 |
| DAPI | Invitrogen | D3571 | NA | 0.5 |
| Anti syntaxin-2 | ProteinTech | 55033-1-AP | NA | 10 |
| Anti GFP | Roche | 11814460001 | 0.4 | 0.8 |
| Anti GRASP65 | Santa Cruz | sc-19481 | NA | 1 |
| Anti Lamp-1 | Santa Cruz | sc-18821 | NA | 0.02 |
| Anti beta-tubulin | Sigma-Aldrich | T4026 | 1:5000 | NA |
| Anti Munc18-1 | Synaptic Systems | 116 003 | 1 | NA |
| Anti syntaxin 1A | Synaptic Systems | 110 118 | NA | 10 |
| Anti syntaxin-2 | Synaptic Systems | 110 123 | 1 | NA |

### Confocal microscopy

High resolution images were acquired in an inverted Leica TCS SP8 microscope equipped with 405, 458, 476, 488, 496, 514, 561, and 633 nm laser lines, photomultipliers (PMT) and hybrid detectors. Unless otherwise indicated, a plan apo 63X, 1.4 NA, oil-immersion objective was used for specimen imaging. Pixel size and z-step sizes were set to fulfil Nyquist criterion. In all cases, laser and detection spectral bands were chosen to maximize signal recovery while avoiding signal bleed-through. Scanning speed was set to 600 Hz and bidirectional. Between 1 and 3 lines were averaged to generate the final image. When sequential scanning was needed to avoid signal bleed-through, each acquisition sequence was done in between lines. Bit-depth was set to 8 bits.

### Live cell imaging

Image acquisition was done in an Andor Revolution XD Spinning Disk confocal microscope equipped with 405, 445, 488, 514, and 561 nm laser lines. Digitalization of the fluorescent emission was done with an EM CCD camera of 512 × 512 pixels and 0.37594 µm xy resolution. Electron multiplier gain was set to 300. Exposure time was set to 300 ms and bit-depth was 14 bits. Imaging medium was 1.8 ml of isotonic buffer (see secretion assays of materials and methods for composition). One complete z-stack acquisition was done every 20 s for 5 minutes before ATP addition. After 5 minutes, 0.2 ml of 100 mM ATP was added to the cells to reach a final ATP concentration of 100 µM. Cells were imaged every 30 s for another 25 minutes.

### Dot and western blot quantification

ImageJ / Fiji (49, 50)were used for the analysis of all digitalized images from dot and western blots experiments. To quantify ECL or fluorescence intensity, the background was subtracted from the images. We generated an intensity profile for the membrane and measured the area of the peaks of interest. These area values are a direct measure of the emission intensity of the bands of interest in the membrane. Only images with no saturated pixels were used for quantification.

### Co-localization quantification

For co-localization analysis we calculated the Pearson Correlation Coefficient (PCC) of confocal images of cultured HT29 cells using Fiji/ImageJ (49, 50). In monocultures, we calculated PCC with the coloc2 plugin included in Fiji/ImageJ. To calculate the PCC for each optical slide of confocal images of the Caco2/HT29 co-cultures we wrote an ImageJ macro to subtract the image’s background and then calculate the PCC of the entire optical slide.

### Quantification of the number of mucin granules in live cell imaging experiments

Quantification was done as previously described (45). Spinning disk confocal images were analysed with the ‘‘spot detector’’ (51) plug-in in Icy software (52). Spots were detected by first processing the original images to obtain coefficient images to remove background and noise. Then, a wavelet adaptive threshold was computed using a combination of scale 2 and 3 with a threshold between 30 and 80 to define the size of the spots and sensitivity of the algorithms respectively. In R, the total number of spots (mucin granules) for each time point (frame) of the time-lapse were calculated by the adding all the detected spots in each optical slide. Finally, the number of spots in each frame were divided by the number of spots in the first frame.

### Statistical analysis

All statistical analysis and graphical representation of the data was performed with the R software, several packages from the tidyerse (53) and the data.table package. When the experimental design required a single pairwise comparison, Student t tests were applied.

When the experimental design required the comparison of multiple conditions with a single control condition an ANOVA with orthogonal contrasts was applied. In the cases where we needed to do multiple comparisons, we applied an ANOVA test followed by a Tukey’s HSD test. When the experimental design contained more than one principal factor, we used two-way ANOVA with interaction. The exact value of n, what it represents and centre and dispersion measures are described in the figures and figure legends. The exact p value of the statistical test is written in the figures or figure legends. Secretion assays were performed by triplicates each time. In the statistical model these triplicates are nested together. The lower and upper hinges of the box plots correspond to the first and third quartiles respectively (the 25th and 75th percentiles). The upper whisker extends from the hinge to the largest value no further than 1.5 * IQR from the hinge (where IQR is the interquartile range, or distance between the first and third quartiles). The lower whisker extends from the hinge to the smallest value at most 1.5 * IQR of the hinge. Data beyond the end of the whiskers are called ‘‘outlying’’ points and are plotted individually.

## Acknowledgments

We thank all members of the Malhotra laboratory, specially Ishier Raote, and Meir Aridor from the University of Pittsburgh School of Medicine for valuable discussions and critical reading of the manuscript. Cell sorting experiments were carried out by the joint CRG/UPF FACS Unit at Parc de Recerca Biomèdica de Barcelona (PRBB). Fluorescence microscopy was performed at the Advanced Light Microscopy Unit at the CRG, Barcelona. We acknowledge the support of the Spanish Ministry of Science and Innovation to the EMBL partnership, the Centro de Excelencia Severo Ochoa, and the CERCA Programme / Generalitat de Catalunya. V. Malhotra is an Institució Catalana de Recerca i Estudis Avançats professor at the Centre for Genomic Regulation. **Funding**. Work in the Malhotra lab is funded by grants from the Spanish Ministry of Economy and Competitiveness (Plan Nacional to VM: PID2019-105518GB-I00) and the European Research Council Synergy Grant (ERC-2020-SyG -Proposal No. 951146). JW is funded by the European Research Council (H2020-MSCA-IF-2019-894115). This work reflects only the authors’ views, and the EU Community is not liable for any use that may be made of the information contained therein.

## Author contributions

Conceptualization, J.W., and V.M.; Methodology, J.W., A.L., G.B., O.F., C.A., M.P.R., and N.B.; Investigation, J.W., A.L., G.B., and V.M.; Data Analysis, J.W.; Writing – Original Draft, J.W. and V.M.; Funding Acquisition, V.M., and J.W.; Supervision, J.W., and V.M.

