## Supplementary Figures for "Biphasic release propensity of mucin granules is supervised by TSPAN8"

**TSPAN8 sequesters syntaxin 2 to prevent docking of mucin filled vesicles thereby keeping them in reserve to control quantities of mucins secreted.**

Wojnacki José[1], Lujan Agustín[1], Foresti Ombretta[1], Aranda Carla[1], Bigliani Gonzalo[1], Pena Rodriguez Maria[1], Brouwers Nathalie[1], and Malhotra Vivek[1;2,3,] .

[1] Centre for Genomic Regulation (CRG), The Barcelona Institute for Science and Technology, Dr. Aiguader 88, 08003 Barcelona, Spain.

[2] Universitat Pompeu Fabra (UPF), Barcelona 08002, Spain

[3] ICREA, Barcelona 08010, Spain

### Supplementary figures

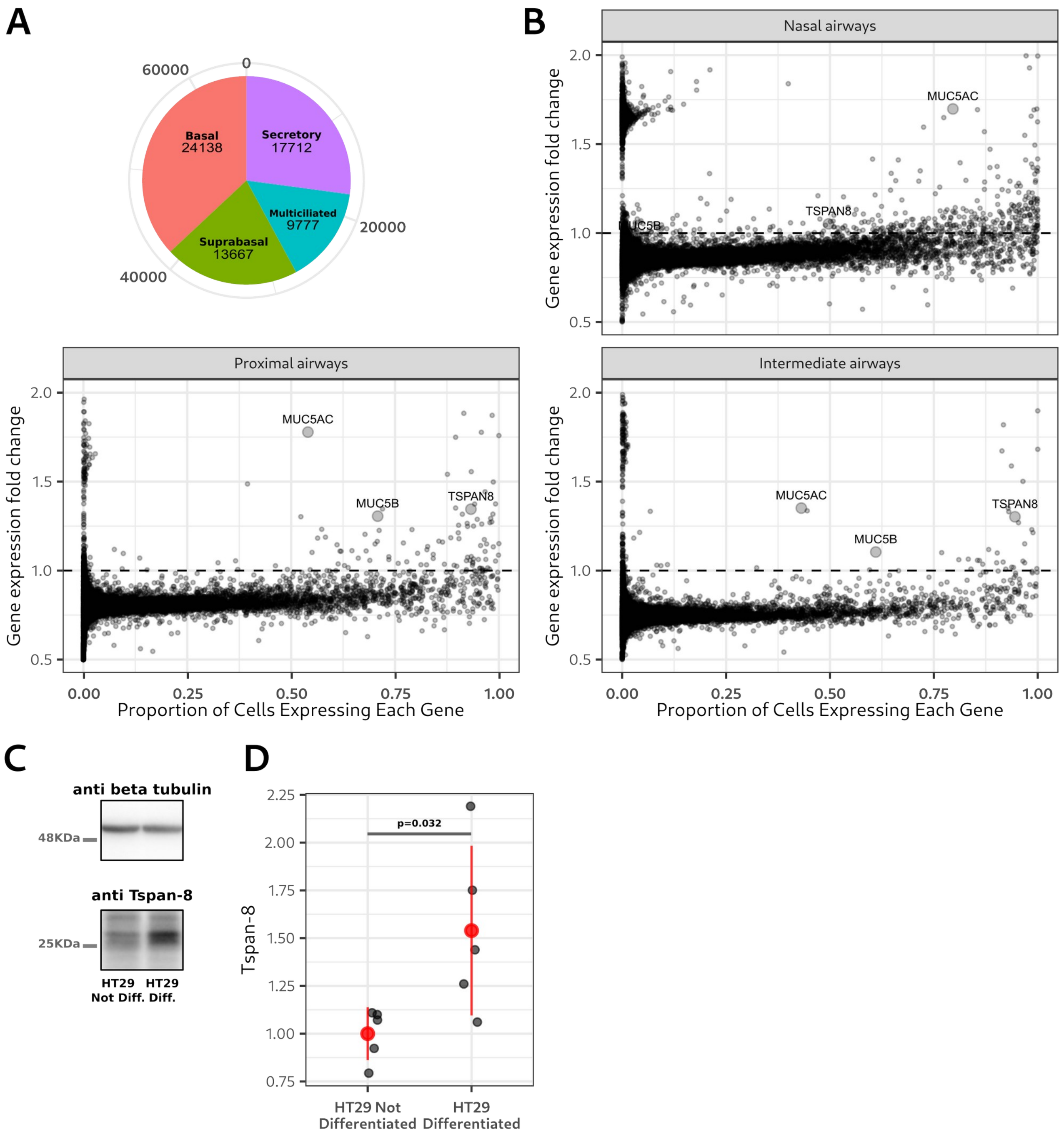


**Supplementary Figure 1. Tetraspanin-8 enrichment in mucin-secreting cells from the healthy human airways.**

**A.** Pie chart showing the total number and type of cells analyzed in the bioinformatics analysis.

**B.** Dot plots showing the fold-change in gene expression between undifferentiated basal and mucin-secreting cells in the y-axis and the proportion of mucin-secreting cells expressing each gene in the xaxis. Each dot represents one gene. Larger dots represent MUC5AC, MUC5B and TSPAN8. Genes below the dashed horizontal line are down regulated in mucin secreting cells while genes above the line are up regulated. Dots were plotted with a 50% transparency value for better visibility in areas of the graph with high dots density.

**C.** Representative western blot of the total amount of beta-tubulin (upper panel) and Tspan-8 (lower panel) detected in growing (non-differentiated) and differentiated HT29 (mucin-secreting) cells. Blots were incubated with anti beta-tubulin and anti Tspan-8 antibodies and developed by enhanced chemiluminescence (ECL).

**D.** Quantification of the total amount of beta-tubulin and Tspan-8 in growing (non-differentiated) and differentiated mucin secreting HT29 cells. A representative membrane is shown in **C.** Each dot represents the signal of an independent replicate. Total number of replicates are 5. Red dots represent the mean +/- the standard deviation. The p value is from a one-way ANOVA analysis.

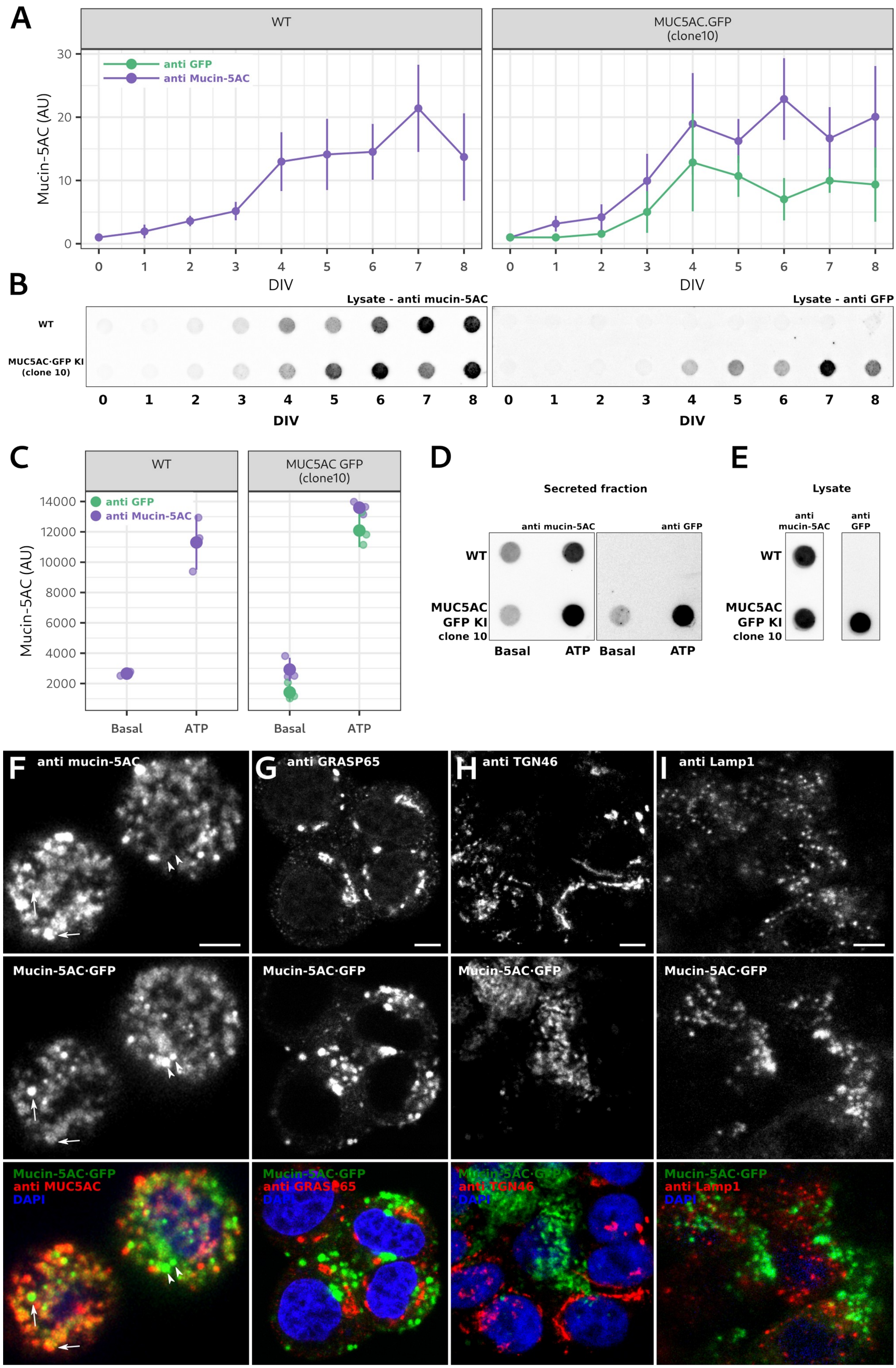
A
WT
MUC5AC.GFP
(clone10)
anti GFP
anti Mucin-5AC
Mucin-5AC (AU)
DIV
B
Lysate - anti mucin-5AC
Lysate - anti GFP
WT
MUC5AC·GFP KI
(clone 10)
DIV
C
WT
MUC5AC GFP
(clone10)
anti GFP
anti Mucin-5AC
Mucin-5AC (AU)
Basal
ATP
D
Secreted fraction
anti mucin-5AC
anti GFP
WT
MUC5AC
GFP KI
clone 10
Basal
ATP
E
Lysate
anti
mucin-5AC
anti
GFP
F
anti mucin-5AC
G
anti GRASP65
H
anti TGN46
I
anti Lamp1
Mucin-5AC·GFP
anti MUC5AC
anti GRASP65
anti TGN46
anti Lamp1
DAPI

**Supplementary Figure 2. Characterization of the MUC5AC CRISPR/Cas9 sfGFP KI cell line.**

**A.** Quantification of the total amount of mucin-5AC during HT29 cell line differentiation as detected by immunolabeling of mucin-5AC (purple line and dots) and GFP (green lines and dots). Representative dot blots membranes are shown in **B** (below). The left and right plots are the quantification of mucin5-AC in WT and mucin-5AC·GFP (identified as clone 10) KI cells respectively. Each dot represents the mean mucin-5AC value of three independent experiments +/- the standard deviation. For each replicate, mucin-5AC values of cells in culture for 1 to 8 days *in vitro* (DIV) were relativized to the value of non-differentiated cells (0 DIV).

**B.** Representative dot blots showing the total amount (cell lysates) of mucin-5AC in HT29 cells during the differentiation process. The left western blot membrane was incubated with an anti mucin-5AC antibody. The right membrane was loaded with identical samples to the left membrane and was incubated with an anti GFP antibody. Signals were detected by fluorescence emission. Top rows are samples from WT cells and the bottom rows are samples from the mucin-5AC·GFP (clone 10) cell line.

**C.** Quantification of the amount of secreted mucin-5AC by unstimulated (basal) and ATP-stimulated (ATP) WT and MUC5AC sfGFP KI (clone 10) cells. A representative dot blot is shown in **D**. Purple dots represent the signal of anti mucin-5AC immunoblotting. Green dots represent the signal from anti GFP immunoblotting. Smaller dots represent the mucin-5AC amount detected from independent secretion assays. Total number of replicates are 3. All samples were processed in parallel. Bigger dots represent the mean secreted mucin-5AC +/- the standard deviation. Signals were detected by fluorescence emission. The left and right plots are the quantification of WT and mucin-5AC·GFP (clone 10) cells, respectively. A two-factor ANOVA analysis showed no statistical difference between WT and mucin-5AC·GFP cell lines ($p = 0.696$) and no difference between anti mucin-5AC and sfGFP signals ($p = 0.646$).

**D.** Representative dot blots of a mucin secretion assay of unstimulated (basal) and ATP-stimulated (ATP) WT and MUC5AC sfGFP KI (clone 10) cells. The left membrane was immunoblotted with an anti mucin-5AC antibody. The right membrane was loaded with identical samples to the left membrane and was immunoblotted with an anti GFP antibody. Signals were detected by fluorescence emission. Top rows are samples from WT cells and the bottom rows are samples from the mucin-5AC·GFP-expressing (clone 10) cell line. For each membrane the left columns are samples from non-stimulated cells (basal) and the right columns are samples of ATP-stimulated cells (ATP).

**E.** Representative dot blots showing the total amount (cell lysate) of mucin-5AC in differentiated WT and mucin-5AC·GFP-expressing (clone 10) cells. The left membrane was immunoblotted with an anti mucin-5AC antibody. The right membrane was loaded with identical samples as in the left membrane and was immunoblotted with an anti GFP antibody. Signals were detected by fluorescence emission. Top row are samples from WT cells and the bottom row are samples from the mucin-5AC·GFP-expressing (clone 10) cell line.

**F – I**. Top row: Optical planes from confocal images of differentiated mucin-5AC·GFP (clone 10) cells and immunolabeled for mucin-5AC (F), GRASP65 (G), TGN46 (H) and Lamp1 (I). Scale bar is 10 µm. Middle row: Images showing mucin-5AC·GFP localization in the same optical plane as in the top row. The GFP signal was not enhanced by antibody detection. Arrows in **F** point to co-localizing anti mucin-5AC and mucin-5AC·GFP signals. Arrowheads point to mucin-5AC·GFP-positive but anti mucin-5AC-negative granules. Bottom row: Merge of images from top and middle rows. DAPI was used to visualize the cell nucleus.

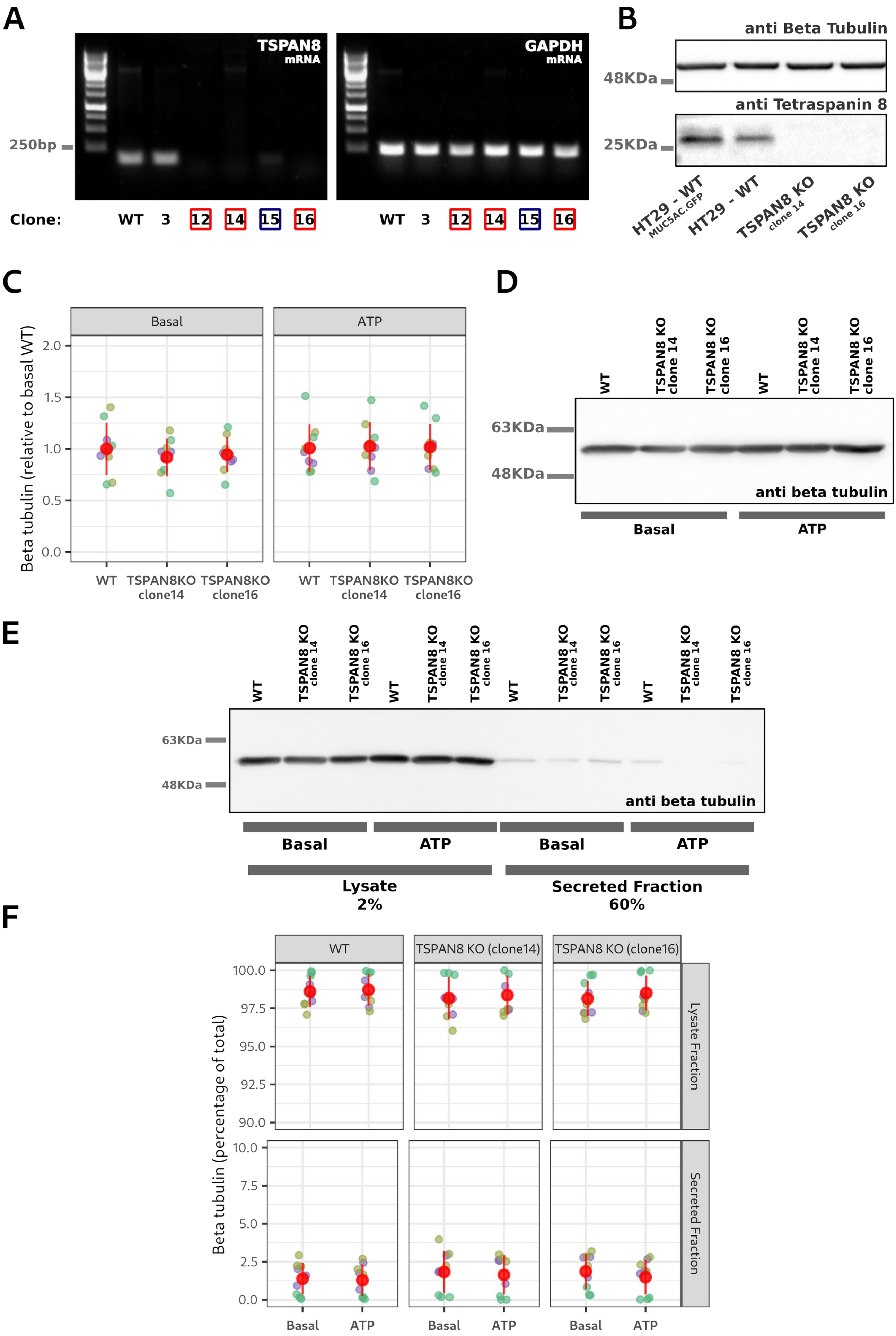

A
TSPAN8 mRNA
GAPDH mRNA
250bp
Clone: WT 3 12 14 15 16
WT 3 12 14 15 16
B
anti Beta Tubulin
48KDa
anti Tetraspanin 8
25KDa
HT29 - WT MUC5AC.GFP
HT29 - WT
TSPAN8 KO clone 14
TSPAN8 KO clone 16
C
Basal
ATP
Beta tubulin (relative to basal WT)
2.0
1.5
1.0
0.5
0.0
WT
TSPAN8KO clone14
TSPAN8KO clone16
D
WT
TSPAN8 KO clone 14
TSPAN8 KO clone 16
63KDa
48KDa
anti beta tubulin
Basal
ATP
E
63KDa
48KDa
anti beta tubulin
Basal
ATP
Lysate 2%
Secreted Fraction 60%
F
WT
TSPAN8 KO (clone14)
TSPAN8 KO (clone16)
Beta tubulin (percentage of total)
100.0
97.5
95.0
92.5
90.0
10.0
7.5
5.0
2.5
0.0
Lysate Fraction
Secreted Fraction
Basal
ATP

**Supplementary Figure 3. Tetraspanin 8 KO generation and secretion assays.**

**A.** CRISPR/Cas9-genetically modified cell lines identified as clones 3, 12, 14, 15 and 16 were screened for the presence of TSPAN8 mRNA. Agarose gels showing TSPAN8 (left) and GAPDH (right) cDNA amplification. cDNA was obtained by RT-PCR from the mRNA purified from the screened cell lines. Red boxes show full TSPAN8 KO and the purple box shows a probable heterozygote cell line. GAPDH was used as a positive RT-PCR control.

**B.** Representative western blot showing the total amount of Tspan-8 (bottom panel) in WT and TSPAN8 KO (clones 14 and 16) cell lines. Beta-tubulin (top panel) was used as loading control. Membranes were immunoblotted with anti Tspan-8 and anti beta-tubulin antibodies and developed by ECL.

**C.** Quantification of the total (cell lysates) beta-tubulin content from the secretion assays quantified in Figure **2B**. A representative western blot is shown in **D**. Each dot represents beta-tubulin signal from an independent secretion assay. Grouped in different colors are secretion assays that were processed in parallel. Total number of replicates are 9. Red dots represent the mean +/- the standard deviation. Values are expressed as relative to the average beta-tubulin content of non-stimulated WT cells. A two-way ANOVA with interaction was done, and no statistical differences were found. Genotype:Secretion interaction p value = 0.929; Secretion principal factor p value = 0.987; Genotype principal factor p value = 0.847.

**D.** Loading control of the dot blot shown in Figure **2A**. Representative western blot of the total amount (cell lysate) of beta-tubulin. The membrane was immunoblotted with an anti beta-tubulin antibody and developed by ECL.

**E.** Representative western blot of the amount of beta-tubulin present in the cell lysates and in the secreted fractions of a secretion assay quantified in Figure **2B**. 2% of the cell lysate and 60% of the secreted fraction were loaded into the gel. The secreted fraction was precipitated with trichloro acetic acid to reduce the total volume of the sample. The membrane was immunoblotted with an anti beta-tubulin antibody and developed by ECL.

**F.** Quantification of the amount of beta-tubulin present in cell lysates and in the secreted fractions of a secretion assay quantified in Figure **2B**. A representative western blot is shown in **E**. Each dot represents the beta-tubulin signal from an independent secretion assay. Grouped in different colors are secretion assays that were processed in parallel. Total number of replicates are 9. Red dots represent the mean +/- the standard deviation. Values are expressed as percentage of the total (lysate and secreted) amount of beta-tubulin. Top panels are the percentage of tubulin in the lysate and lower panels are beta-tubulin in the secreted fractions. A two-way ANOVA with interaction was done, and no statistical differences were found. Genotype:Secretion interaction p value = 0.940; Secretion principal factor p value = 0.488; Genotype principal factor p value = 0.556.

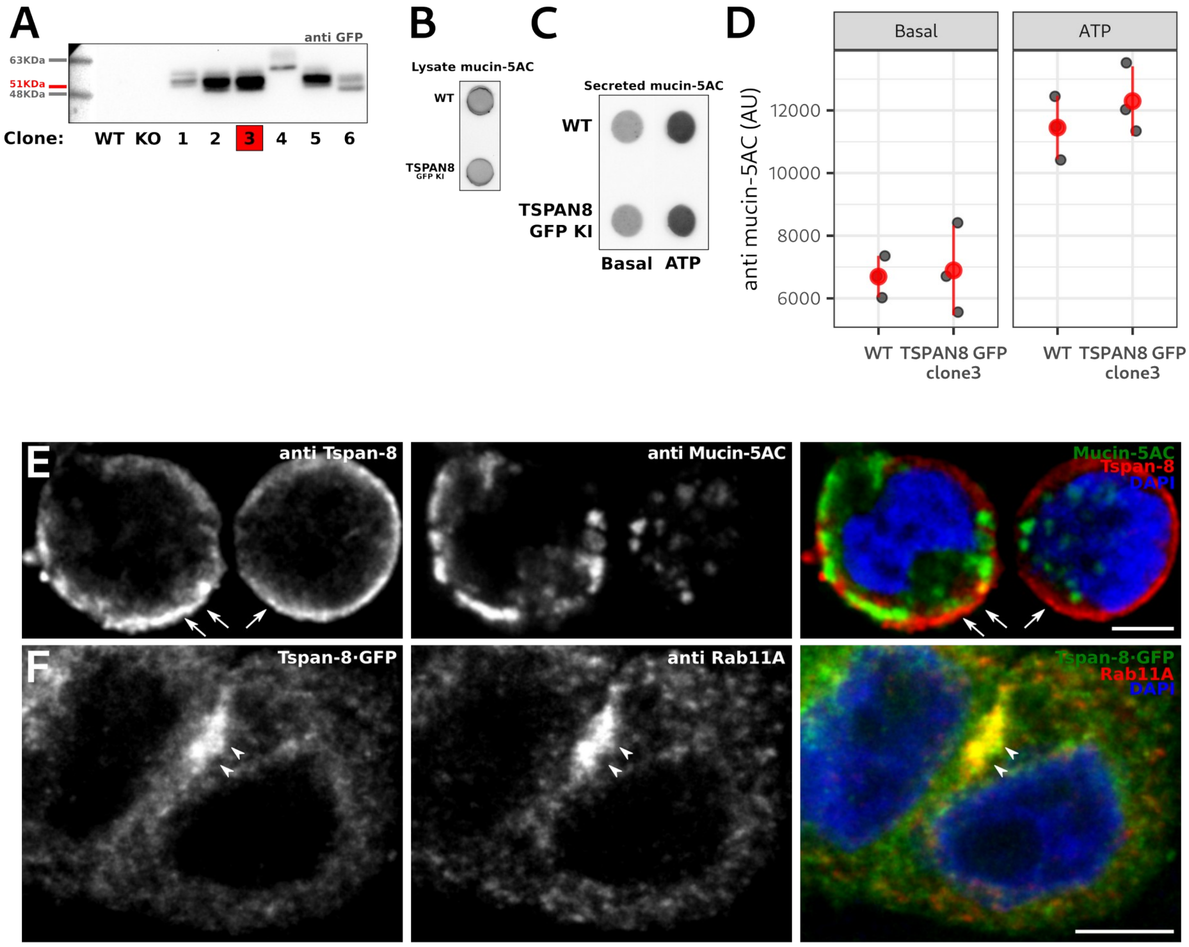


**Supplementary Figure 4. Generation of a tetraspanin 8·GFP-expressing cell line by CRISPr/Cas.**

**A.** Western blot screening of 6 different cell lines modified to express sfGFP at the c-terminus of TSPAN8. Lysates from WT and TSPAN8 KO cells were used as control. Lines labeled 1 to 6 are the cell lysates of 6 different clones screened for the presence of Tspan-8·GFP. The membrane was immunoblotted with an anti GFP antibody and developed by ECL. The red tick shows the expected migration pattern of the fusion protein Tspan8·GFP (26KDa from Tspan-8 and 25KDa from sfGFP). Clone 3 was used for all subsequent experiments.

**B.** Representative dot blot showing the mucin-5AC content of WT and Tspan-8·GFP (clone 3) cell lysates. The membrane was immunoblotted with an anti mucin-5AC antibody. The signal was detected by fluorescence emission.

**C**. Representative dot blot of a secretion assay showing mucin-5AC content in the secreted fractions of WT and Tspan-8·GFP (clone 3) in non-stimulated (basal) and ATP-stimulated cell lines. The membrane was immunoblotted with an anti mucin-5AC antibody. The signal was detected by fluorescence emission.

**D.** Quantification of the secretion assays of non-stimulated (basal) or ATP-stimulated WT and Tspan-8·GFP (clone 3) cell lines. A representative dot blot is shown in **C**. Each gray dot represents the mucin-5AC amount of an independent secretion assay. All three secretion assays were run in parallel. The red dot is the mean value of the gray dots +/- the standard deviation. The y-axis is in arbitrary units (AU). A two-way ANOVA with interaction of the secreted mucin-AC was done. Genotype:Secretion interaction p value = 0.622; Secretion principal factor (basal vs ATP) p value < 0.001; Genotype principal factor p value = 0.431.

**E.** Optical plane from a confocal image of WT mucin-secreting cells immunolabelled for Tspan-8 (left image) and mucin-5AC (middle image). Right image is the merge of the two. DAPI was used to visualize the cell nucleus. Arrows point to a region of the cell that is not proximal to the cell nucleus. Scale bar is 10 μm.

**F.** Optical image of a representative confocal image of a mucin-secreting cell expressing Tspan-8·GFP (left image) at endogenous levels and immunolabelled against Rab11A (middle image). Right image shows the merge of the two. DAPI was used to visualize the cell nucleus. The GFP signal was enhanced by antibody detection of GFP. Arrowheads point to recycling endosomes as determined by its localization and Rab11A signal. Scale bar is 10 μm.

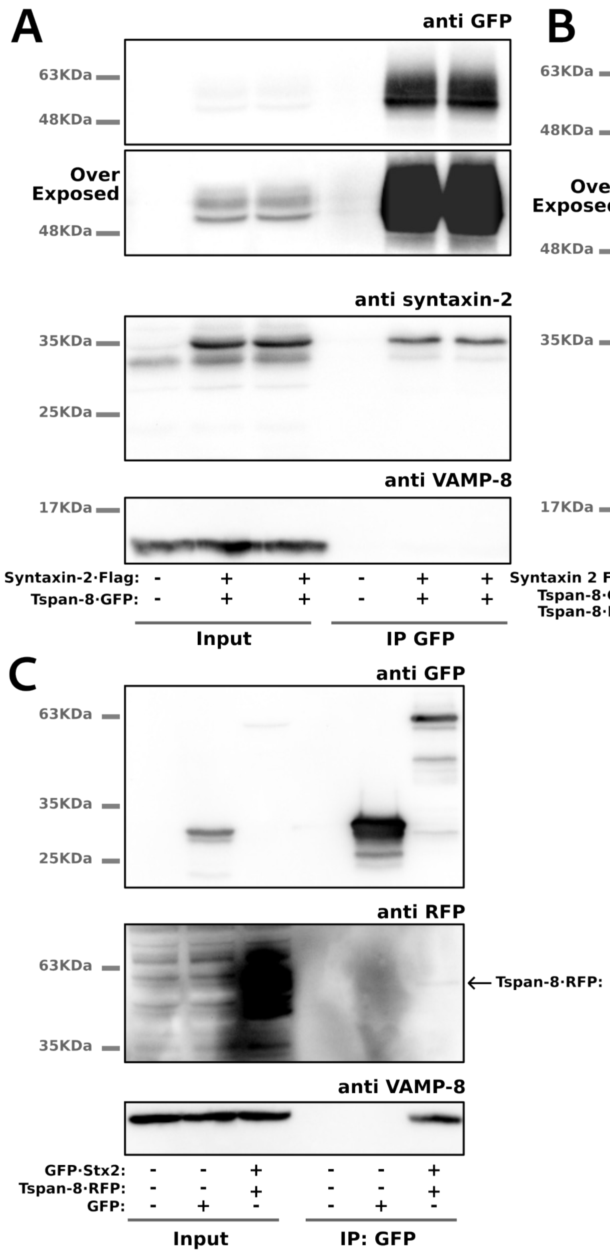


**Supplementary Figure 5. Tspan-8 interacts with syntaxin 2.**

**A – B.** Lysates of WT HT29 cells transiently co-transfected with Tspan-8·GFP and Stx2·FLAG were processed for GFP immunoprecipitation and western blot analysis. Top panels show immunoblotting against GFP to confirm the immunoprecipitation. Middle-top panels show the over-exposed GFP-immunoblotted membranes. Middle-lower panels show the immunoblotting against Stx2. Lower panels show immunoblotting against VAMP-8. No transfection (**A** and **B**) and Tspan-8·RFP transfection (**B**) were used as control conditions. In A, 2 independent test conditions are shown in the same membrane.

**C.** Lysates of WT HT29 cells transiently co-transfected with GFP·Stx2 and Tspan-8·RFP were processed for GFP immunoprecipitation and western blot analysis. Top panel shows immunoblotting against GFP to confirm the immunoprecipitation. Middle panel shows the immunoblotting against RFP. Lower panel shows immunoblotting against VAMP-8. No transfection and transfection of soluble GFP were used as control conditions. The arrow points to the Tspan-8·RFP band on the co-immuno-precipitate.

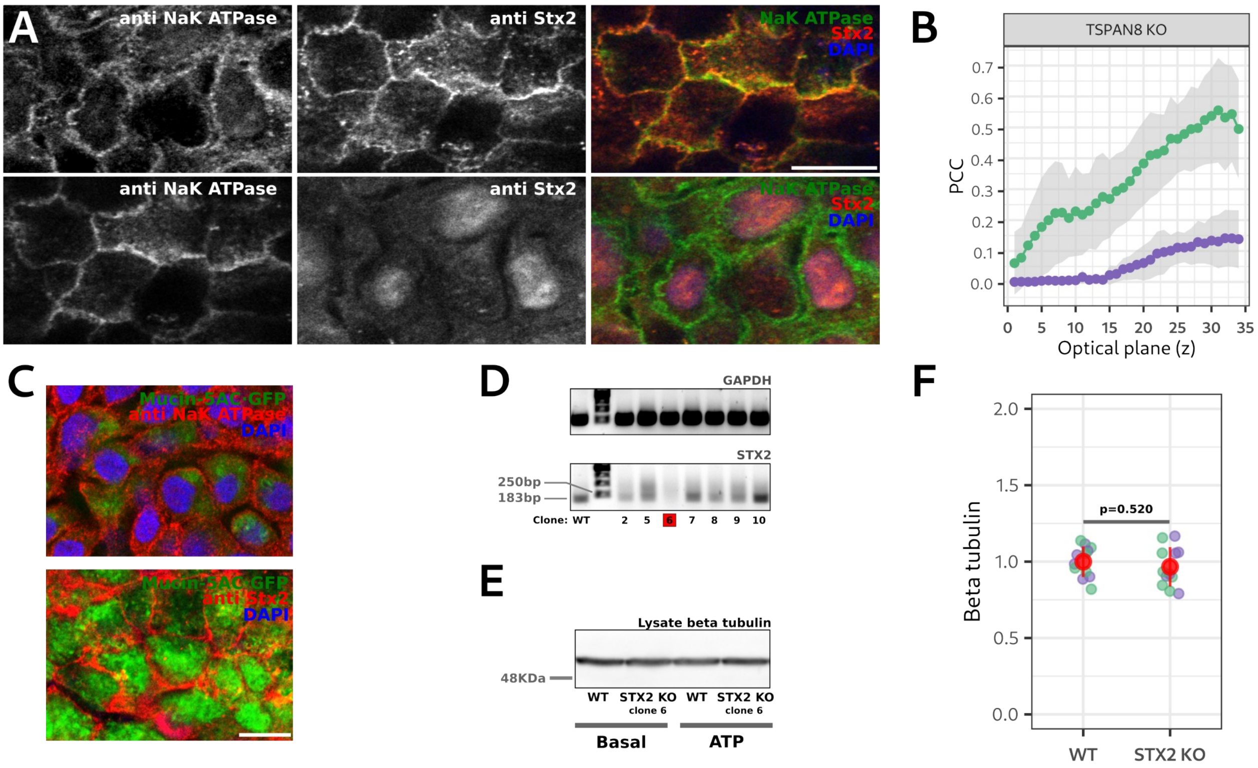


**Supplementary Figure 6. Syntaxin-2 in necessary for mucin-5AC secretion.**

**A.** Optical planes of a representative confocal image of a HT29 TSPAN8 KO/Caco2 co-culture immunolabelled for $Na^+/K^+$-ATPase (left images) and Stx2 (middle images). Right images show the merge of the two and DAPI to visualize the cell nucleus. Scale bar is 10 µm. Top row is an optical plane of the apical part of the co-culture and the lower row is an optical plane of the basal/medial part of the co-culture and distinguishable by the presence of DAPI staining.

**B.** Green dots and lines show the mean PCC quantification between the $Na^+/K^+$-ATPase and Stx2 signals in each optical plane of HT29 TSPAN8 KO/Caco2 co-cultures. Representative images are shown in **A**. Purple dots and lines show the mean PCC between Stx2 and mucin-5AC·GFP. The gray areas show the mean PCC +/- the standard deviation.

**C**. Optical plane of a representative confocal image of a HT29 WT/Caco2 co-culture expressing mucin-5AC·GFP and immunolabelled for $Na^+/K^+$-ATPase (upper image) and Stx2 (lower image). DAPI was used to visualize the cell nucleus. Scale bar is 10 µm. Left image shows the basal/medial part of the co-culture and is distinguishable by the presence of DAPI staining and little, if any, mucin-5AC·GFP. The right image shows an optical plane of the apical part of the co-culture and is distinguishable by the high abundance of mucin-5AC·GFP and the lack of DAPI staining.

**D.** Grown cell lines identified as clones 2, 5, 6, 7, 8, 9 and 10 were screened for the presence of STX2 mRNA after CRISPR/Cas9 genome editing. Agarose gels showing STX2 (lower panel) and GAPDH (top panel) cDNA amplification. cDNA was obtained by RT-PCR from the mRNA purified from the screened cell lines. Red box shows a full STX2 KO clone. GAPDH was used as a positive RT-PCR control. Clone 6 was used for subsequent experiments.

**E**. Representative western blot of the total amount (cell lysate) of beta-tubulin from the secretion assay shown in Figure **5D**. The membrane was immunoblotted with an anti beta-tubulin antibody and developed by ECL.

**F.** Quantification of the total (cell lysates) beta-tubulin content from the samples of the secretion assays quantified in Figure **5E**. A representative western blot is shown in **E**. Each dot represents beta-tubulin signal from a single secretion assay. Grouped in different colors are samples from basal and ATP-stimulated cells. Total number of replicates are 9. Red dots represent the mean +/- the standard deviation. Values are expressed as relative to the average beta-tubulin content in WT cells. A two-way ANOVA with interaction was done, and no statistical differences were found. Genotype:Secretion interaction p value = 0.757; Secretion principal factor p value = 0.742; Genotype principal factor p value = 0.520.

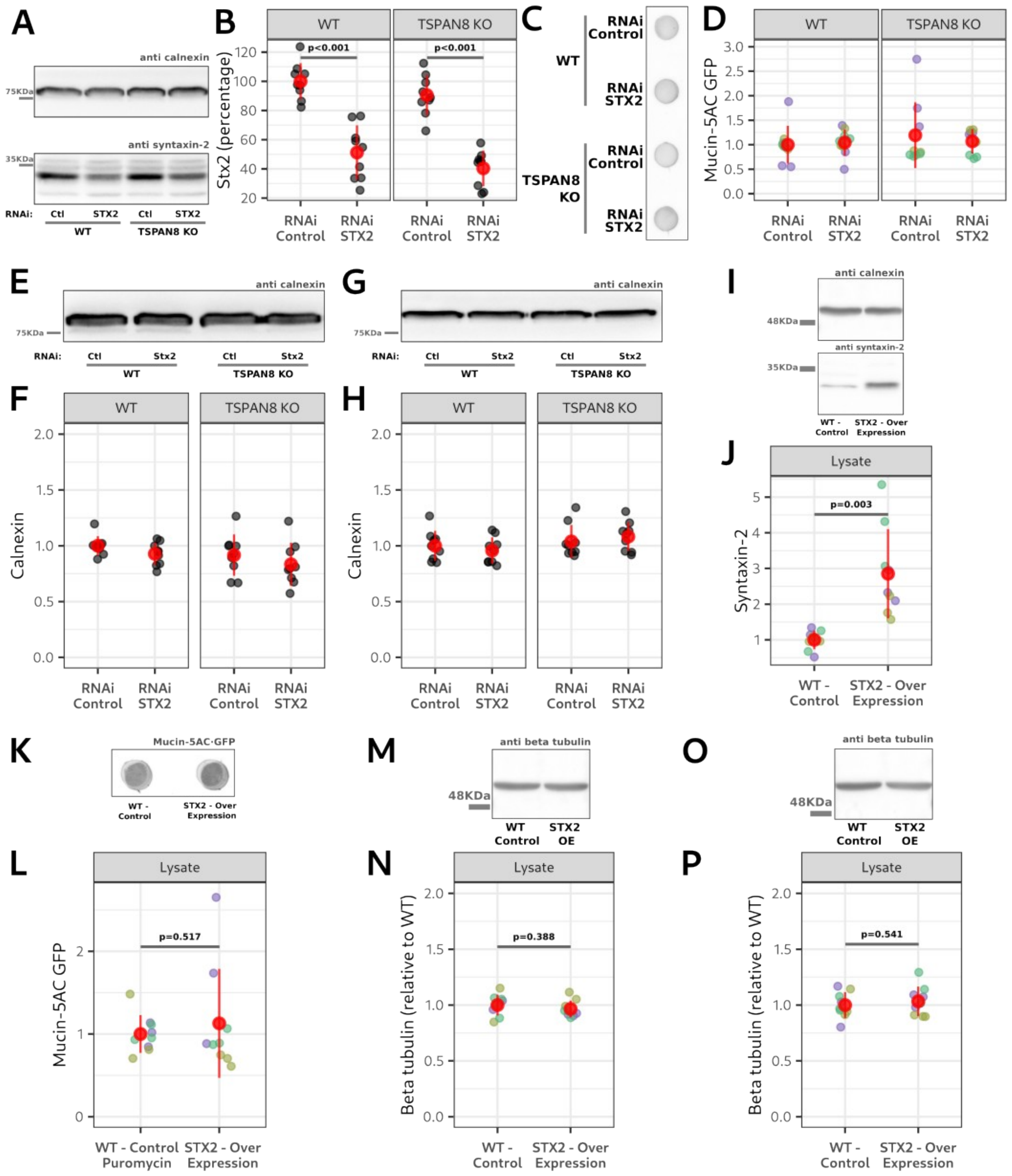


**Supplementary Figure 7. Syntaxin-2 and Tspan-8 are in the same pathway of regulated mucin secretion.**

**A.** Representative western blot of the total amount (cell lysates) of Stx2 (lower panel) and calnexin (upper panel) present in WT and TSPAN8 KO cells treated with RNAi control and RNAi against STX2. Membranes were immunoblotted with anti calnexin and anti Stx2 antibodies, respectively and developed by ECL.

**B.** Quantification of the amount of Stx2 detected in WT and TSPAN8 KO cells treated with RNAi control and RNAi against STX2. A representative western blot is shown in **A**. Each dot represents Stx2 signal from an independent sample. Total number of replicates are 9. Stx2 signals were divided by the corresponding calnexin signals to control for equal sample loading in the western blot. Red dots represent the mean +/- the standard deviation. Values are expressed as percentages. 100% is the mean total amount of Stx2 in WT cells treated with control RNAi. Left panel is the quantification of WT cells and the right panel is the quantification of TSPAN8 KO cells. A two-way ANOVA with interaction of the total Stx2 amount was done. The Genotype:Treatment interaction p value = 0.935 shows that RNAi treatment equally affected WT and TSPAN8 KO cells; Genotype principal factor p value = 0.113; Treatment principal factor p value < 0.001. p values in the graphs are from a TukeyHSD post hoc statistical test.

**C.** Representative dot blot of the total (cell lysate) mucin-5AC·GFP in WT and TSPAN8 KO cells treated with RNAi control and RNAi against STX2. The fluorescent emission of sfGFP is shown.

**D.** Quantification of the amount of mucin-5AC·GFP in the cell lysates of WT and TSPAN8 KO cells treated with RNAi control and RNAi against STX2. A representative dot blot is shown in **C**. Each dot represents signal from an independent sample. Grouped in different colors are samples that were processed in parallel. Total number of replicates are 9. Red dots represent the mean +/- the standard deviation. A two-way ANOVA with interaction was done, and no statistical differences were found. Genotype:Treatment interaction p value = 0.460; Genotype principal factor p value = 0.342; Treatment principal factor p value = 0.723.

**E.** Loading control of the dot blot shown in **C**. Representative western blot of the total amount (cell lysate) of calnexin. The membrane was immunoblotted with an anti calnexin antibody and developed by ECL.

**F.** Quantification of the total (cell lysates) calnexin content from the samples used in the dot blots quantified in **D**. A representative western blot is shown in **E**. Each dot represents calnexin signal from an independent sample. Total number of replicates are 9. Red dots represent the mean +/- the standard deviation. Values are expressed as relative to the average calnexin content in WT control cells. A two-way ANOVA with interaction was done, and no statistical differences were found. Genotype:Treatment interaction p value = 0.926; Treatment principal factor p value = 0.163; Genotype principal factor p value = 0.111.

**G.** Loading control of the secretion assay shown in Figure **6C**. Representative western blot of the total amount (cell lysate) of calnexin. The membrane was immunoblotted with an anti calnexin antibody and developed by ECL.

**H.** Quantification of the total (cell lysates) calnexin content from the secretion assay quantified in Figure **6D**. A representative western blot is shown in **G**. Each dot represents calnexin signal from a single secretion assay. Total number of replicates are 9. Red dots represent the mean +/- the standard deviation. Values are expressed as proportions relative to the average calnexin content in WT control cells. A two-way ANOVA with interaction was done, and no statistical differences were found. Genotype:Treatment interaction p value = 0.371; Treatment principal factor p value = 0.997; Genotype principal factor p value = 0.095.

**I.** Representative western blot of the total amount (cell lysates) of Stx2 (lower panel) and calnexin (upper panel) present in WT and STX2-over expressing cells. Membranes were immunoblotted with anti calnexin and anti Stx2 antibodies and developed by ECL.

**J.** Quantification of the amount of Stx2 detected in WT and STX2-over expressing cells. A representative western blot is shown in **I**. Each dot represents Stx2 signal from an independent sample. Total number of replicates are 9. Grouped in different colors are samples that were processed in parallel. Red dots represent the mean +/- the standard deviation. Stx2 signals were divided by the corresponding calnexin signals to control for equal sample loading in the western blot. Values are expressed as proportions and relative to the mean Stx2 amount in control WT cells. A one-way ANOVA of the total Stx2 amount was done and the p value is shown in the graph.

**K.** Representative dot blot of the total (cell lysate) mucin-5AC·GFP detected in control and STX2-over expressing cells. The fluorescent emission of sfGFP is shown.

**L.** Quantification of the detected amount of mucin-5AC·GFP in the cell lysates of control and STX2-over expressing cells. A representative dot blot is shown in **K**. Each dot represent the signal from a single sample. Grouped in different colors are samples that were processed in parallel. Total number of replicates are 9. Red dots represent the mean +/- the standard deviation. A one-way ANOVA was done and the p value is shown in the graph.

**M.** Loading control of the dot blot shown in **K**. Representative western blot of the total amount (cell lysate) of beta-tubulin. The membrane was immunoblotted with an anti beta-tubulin antibody and developed by ECL. OE refers to over expression.

**N.** Quantification of the total (cell lysates) beta-tubulin content from the samples used in the dot blots quantified in **L**. A representative western blot is shown in **M**. Each dot represents the beta-tubulin signal from a single sample. Grouped in different colors are samples that were processed in parallel. Total number of replicates are 9. Red dots represent the mean +/- the standard deviation. Values are expressed as relative to the average beta-tubulin content in WT control cells. A one-way ANOVA was done and the p value is shown in the graph.

**O.** Loading control of the dot blot shown in Figure **6E**. Representative western blot of the total amount (cell lysate) of beta-tubulin. The membrane was immunoblotted with an anti beta-tubulin antibody and developed by ECL. OE refers to over expression.

**P.** Quantification of the total (cell lysates) beta-tubulin content from the secretion assay quantified in Figure **6F**. A representative western blot is shown in **O**. Each dot represents the beta-tubulin signal from an independent secretion assay. Grouped in different colors are secretion assays that were run in parallel. Total number of replicates are 9. Red dots represent the mean +/- the standard deviation. Values are expressed as relative to the average beta-tubulin content in WT control cells. A one-way ANOVA was done and the p value is shown in the graph.
